# SARS-CoV-2 spike protein as a bacterial lipopolysaccharide delivery system in an overzealous inflammatory cascade

**DOI:** 10.1101/2021.10.29.466401

**Authors:** Firdaus Samsudin, Palur Raghuvamsi, Ganna Petruk, Manoj Puthia, Jitka Petrlova, Paul MacAry, Ganesh S. Anand, Artur Schmidtchen, Peter J. Bond

**Affiliations:** Bioinformatics Institute (BII), Agency for Science, Technology and Research (A*STAR), Singapore 138671, Singapore; Department of Biological Sciences, National University of Singapore, Singapore 117543, Singapore; Division of Dermatology and Venereology, Department of Clinical Sciences, Lund University, SE-22184 Lund, Sweden; Xinnate AB, Medicon Village, SE-22381 Lund, Sweden; Life Sciences Institute, Centre for Life Sciences, National University of Singapore, 28 Medical Drive, Singapore 117546, Singapore; Department of Chemistry, The Pennsylvania State University, 104 Chemistry Building, University Park, PA, USA; Copenhagen Wound Healing Center, Bispebjerg Hospital, Department of Biomedical Sciences, University of Copenhagen, DK-2400 Copenhagen, Denmark

## Abstract

Accumulating evidence indicates a potential role for bacterial lipopolysaccharide (LPS) in the overactivation of the immune response during SARS-CoV-2 infection. LPS is recognised by Toll-like receptor 4 (TLR4) in innate immunity. Here, we showed that LPS binds to multiple hydrophobic pockets spanning both the S1 and S2 subunits of the SARS-CoV-2 spike (S) protein. LPS binds to the S2 pocket with a lower affinity compared to S1, suggesting its possible role as an intermediate in the TLR4 cascade. Congruently, nuclear factor-kappa B (NF-κB) activation *in vitro* is strongly boosted by S2. *In vivo*, however, a boosting effect is observed for both S1 and S2, with the former potentially facilitated by proteolysis. Collectively, our study suggests the S protein may act as a delivery system for LPS in host innate immune pathways. The LPS binding pockets are highly conserved across different SARS-CoV-2 variants and therefore represent potential therapeutic targets.

## Introduction

Coronavirus disease 2019 (COVID-19) caused by the severe acute respiratory syndrome coronavirus 2 (SARS-CoV-2) has so far infected over 200 million people worldwide and resulted in more than 4 million deaths. While most infections are presented with mild to moderate symptoms, a small number of people develop severe respiratory complications requiring intensive care and mechanical ventilation.^1^ Pulmonary and systemic hyperinflammation are some of the prominent hallmarks of severe COVID-19 disease. These dysregulated inflammatory reactions trigger the onset of sepsis and acute respiratory distress syndrome (ARDS), both of which have been documented in nearly all deceased patients.^1–4^ Sepsis and ARDS are initiated by a full-blown activation of the immune response, typically via pattern recognition receptors such as the Toll-like receptors (TLRs), leading to a cytokine storm that causes excessive and damaging inflammatory effects.^5^ How SARS-CoV-2 infection results in overzealous inflammatory responses remains elusive.

Patients with metabolic syndrome, a cluster of comorbidities including diabetes, hypertension, and obesity, are at a higher risk to develop severe COVID-19 disease involving sepsis and ARDS.^2,6,7^ Interestingly, metabolic syndrome is associated with a high blood level of bacterial endotoxins or lipopolysaccharide (LPS) due to gut dysbiosis and translocation of bacterial components into the systemic circulation.^8,9^ LPS is the main component of the outer membrane of Gram-negative bacteria and is a well characterized pathogen-associated molecular patterns (PAMP) recognised by TLR4^10^ in complex with its lipid binding co-receptor MD-2.^11,12^ LPS contains lipid A, whose archetypal TLR4-stimulatory chemical structure consists of six acyl tails connected to a phosphorylated diglucosamine headgroup (Figure 1A), which is covalently connected to a long and heterogeneous polysaccharide chain. LPS also binds to LPS binding protein (LBP) and to an intermediary receptor, CD14. ^13^ LPS transfer through this series of proteins is a potent trigger of the inflammatory response in sepsis.^14^ COVID-19 patients with cardiac involvement have been shown to have an increased plasma level of LBP, suggesting a degeneration of the gut-blood barrier causing leakage of microbial products.^15^ Synergistic interactions during an acute viral infection between downregulation of angiotensin converting enzyme (ACE2) receptor – the primary cellular receptor of SARS-CoV-2 – with obesity and diabetes have been proposed to result in further impairment of endothelial and gut barrier functions.^16^ A clinical report on severely ill COVID-19 patients with pneumonia demonstrated significantly elevated levels of bacterial LPS.^17^ Non-survivor COVID-19 patients also presented increased systemic level of LPS and soluble CD14 during hospitalisation compared to survivors, indicating a link between elevated LPS level and death.^18^ Using a combination of *in vitro* and *in vivo* experiments, we previously showed that SARS-CoV-2 spike (S) protein can bind to LPS with a similar affinity to CD14, and boosts proinflammatory activity,^19^ thus providing a potential link between high-risk COVID-19 patients and observed symptoms like ARDS and sepsis. Nevertheless, the detailed molecular mechanisms underlying the interactions between the S protein and LPS and how these interactions result in an increased inflammatory response are still unclear.

**Figure 1:**
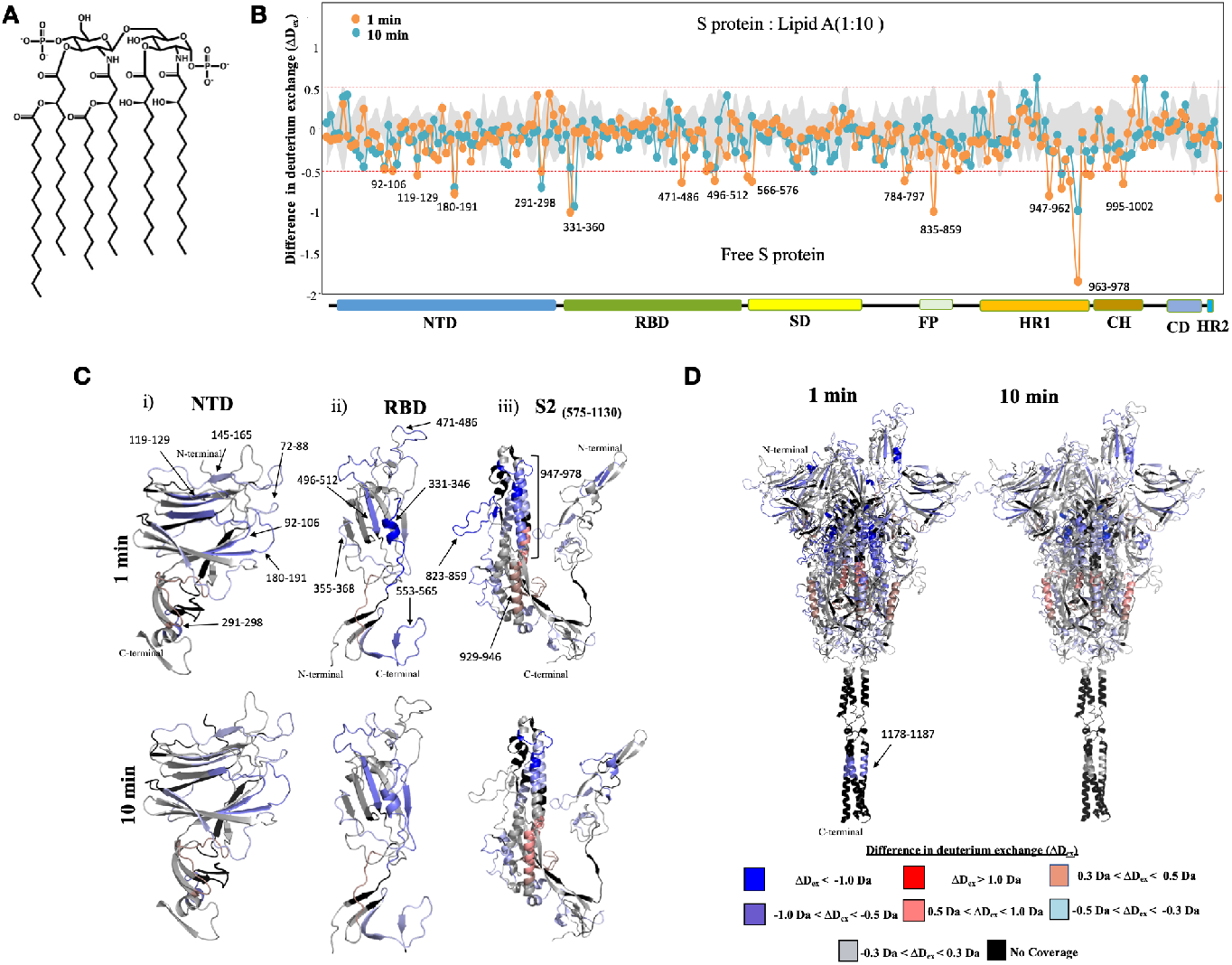
Identifying the orthosteric and allosteric effects of lipid A binding on S protein using HDX-MS. (A) Plot showing differences in deuterium exchange (ΔD_ex_) between S protein: lipid A state and free S protein state at deuterium labelling times t = 1 and 10 min. Pepsin proteolyzed peptide fragments are grouped according to the S protein domain organisation from N to C-termini and plotted along X-axis and its corresponding difference in deuterium exchange values (ΔD_ex_) along Y-axis. A difference cut-off of ±0.5 Da is the significance threshold indicated with red dotted line. (B) Differences in deuterium exchange values at labelling times 1 and 10 min for S protein domains NTD (i), RBD (ii) and S2 domain (iii) are mapped on to the respective structures (C) Differences in deuterium exchange values at labelling times t= 1 min and 10 min are mapped on to the full-length S protein (14-1208) structure. Deuterium exchange differences are colour coded as per key.

The S protein is a class 1 viral fusion protein trimer that protrudes from the surface of SARS-CoV-2.^20,21^ The protein is made of two subunits: the S1 subunit that is involved in receptor binding, and the S2 subunit that facilitates membrane fusion. Proteolysis at the S1/S2 cleavage site by host cell proteases such as furin,^22^ followed by a second proteolytic cleavage on the S2’ site, results in shedding of the S1 subunit and a conformational transition of the S2 subunit into the post-fusion state,^23^ which leads to fusion with the host cell membrane and initiation of infection. The S1 subunit may be further divided into subdomains such as the N-terminal domain (NTD) and the receptor binding domain (RBD), while the S2 subunit is composed of the fusion peptide, heptad repeats, central helix, connector domain and the transmembrane domain. The RBD defines the functional conformation of the S protein, whereby the “open” conformation is characterised by at least one RBD in the “up” conformation which is able to bind to the ACE2 receptor, while the “closed” conformation is characterised by all three RBDs adopting the “down” conformation therefore making them inaccessible to ACE2. The structure of the prefusion S protein ectodomain (ECD) has been resolved in both open and closed conformations using cryo-electron microscopy (cryo-EM)^21,23,24^. Importantly, accumulating structural evidence from cryo-EM has revealed that S protein is able to bind various non-polar molecules such as linoleic acid^25^, polysorbate^26^ and haem metabolites^27^ by employing hydrophobic cryptic pockets in the NTD and RBD. A neutron reflectometry study showed the SARS-CoV-2 S protein significantly degrades lipid bilayers by extracting lipids from the membrane, suggesting a strong association of the protein with lipid molecules^28^. Using computational docking and molecular dynamics (MD) simulations, we predicted that LPS may bind to an inter-protomeric groove nearby the S1/S2 cleavage site.^19^ Similar to the pockets found on the NTD and RBD, this groove has a group of partially exposed hydrophobic residues that could accommodate the LPS lipid tails, primarily found on the S2 subunit (hence designated as the “S2 pocket”). It is not known, however, if LPS can bind to the NTD and RBD pockets.

Structural characterization of LPS binding to proteins using methods such as cryo-EM and X-ray crystallography is challenging due to its heterogenous nature. In this study, we used hydrogen-deuterium exchange mass spectrometry (HDXMS) to verify LPS binding to the S2 pocket of the S protein, while uncovering binding to the NTD and RBD pockets. This was supported by native gel electrophoresis showing LPS interaction with both S1 and S2 subunits independently. Calculation of potentials of mean force (PMFs) within a molecular dynamics (MD) simulation framework, supported by microscale thermophoresis (MST) binding experiments, indicate that LPS binds with strong affinities to the RBD and NTD pockets on the S1 subunit, while the S2 pocket represents a weaker binding site compared to the LPS receptor, CD14, suggesting its potential role as an intermediate in the LPS receptor transfer cascade.^29^ Furthermore, monocytic THP-1 cell assays show boosting of nuclear factor-kappa B (NF-κB) activation by LPS in the presence of the S2 subunit, but not with the S1 subunit. Finally, NF-κB reporter mice experiments show enhanced inflammatory response with both S1 and S2 subunits of the S protein. Collectively, a molecular mechanism of how the SARS-CoV-2 S protein augments LPS-mediated hyperinflammation emerges, whereby the S protein acts as an additional LPS delivery system to the CD14 receptor, and eventually to the TLR4:MD-2 complex.

## Results

### Structural detection of lipid A/LPS binding to SARS-CoV-2 spike protein

To map binding and allosteric effects of lipid A or LPS on S protein, we performed HDXMS on free S protein and S protein saturated with either lipid molecules. Our HDXMS analysis identified pepsin fragment peptides (217 peptides in free S protein state) covering ∼82% of the S protein sequence in the free and bound states (Figure S1). HDXMS provides a readout of solvent accessibility and hydrogen-bonding propensity at a peptide level resolution, and may be used to capture both orthosteric and allosteric effects of ligand/partner protein binding upon protein structure.^30–33^ HDXMS has previously been used to characterize large-scale dynamics of the S protein in the presence of the ACE2 receptor^34^ and an alternative open trimer conformation of the ECD.^35^ Comparative deuterium exchange analysis was performed using deuterium exchange difference plots, wherein the difference in deuterons exchanged for each pepsin proteolyzed peptide at different deuterium labelling times (t =1, 10, and 100 min) between bound and free S protein states were plotted. A difference in deuterium exchange of ± 0.5 Da was chosen as the significance threshold^36^ to identify lipid A/LPS binding induced effects on the S protein.

Deuterium exchange difference plots mapped onto the structure of S protein revealed multiple non-contiguous loci spanning multiple domains – RBD, NTD and S2 domain (Figure 1B-D, S2 and S3) – that were deuterium exchange protected in the lipid A or LPS bound states. Interestingly, protected peptides spanning the RBD (residues 331-360, 496-512) and NTD (residues 92-106, 119-129, 180-191) highlight cavities, optimal for binding lipid A (Figure 1B, 1C(i) and 1C(ii)). These lipid binding cavities were reported in two separate cryo-EM studies wherein densities corresponding to a polysorbate 80 detergent or lipid tails were identified^25,26^. Antiparallel β-strands form a sandwich structure, a convenient binding site for aliphatic tails of lipid A. In the case of the RBD, the lipid A binding cavity consists of a short α-helical region called the “gating helix” along with antiparallel β-strands. Further, several peptides spanning the NTD (residues 291-298), subdomain (SD, residues 538-565), S2 subunit (residues 835-859), and HR1 domain (residues 947-978) show protection from deuterium exchange in the presence of lipid A (Figure 1C(iii)). These peptides are at the inter-protomer interface of the trimeric S protein, forming a hydrophobic cavity which coincides with our previously computationally predicted LPS interaction site.^19^ Interestingly, this region is located close to the functionally significant S1/S2 cleavage site which plays a critical role in viral entry (Figure 1D).

Overall, S protein:lipid A and S protein:LPS bound states show comparable deuterium exchange profiles, and similar peptides were observed to bind lipid A and LPS (Figure 1B-D, S2 and S3). Increases in deuterium exchange were observed in peptides spanning NTD and RBD (residues 72-88, 94-103, 145-165, 178-191, 356-375 and 400-421) at 1 min deuterium labelling time in the presence of LPS (Figure S3A, S3B(i) and S3B(ii)). The increases in deuterium exchange indicate the local destabilization of binding cavities at NTD to accommodate the considerably larger and more heterogeneous LPS molecule compared to lipid A. Peptides spanning the RBD (peptide 331-346, 496-512) and S2 cavity (peptide 835-859, 946-978) show protection at 1 and 10 min labelling times (Figure S3B(ii)-S3B(iii)) similar to the lipid A bound state. Peptide 1178-1187 spanning the HR2 region also showed protection in the presence of both lipid A and LPS at 1 and 10 min labelling times, which is corroborated by recent computational predictions and an NMR study^37,38^ suggesting the potential for binding of hydrophobic molecules such as lipids. An interesting similarity between lipid A and LPS bound S protein is seen with the protection from deuterium exchange at the receptor binding motif (RBM) of the RBD, a crucial component involved in ACE2 receptor interactions.

Interestingly, at longer labelling times (t = 100 min), increases in deuterium exchange were observed across multiple regions on the S protein, especially at the NTD, RBD and S2 binding sites in the presence of lipid A or LPS (Figure S2 and Figure S3). Increases in deuterium exchange at NTD (peptide 32-48, 177-186, 266-277) and RBD (peptide 355-379, 400-421) cavities may result from widening or destabilization of the local structure to accommodate more of the bulky, multiply-acylated lipid A component of LPS, in accordance with our simulation data presented below. Long-range effects of binding to these two pockets have also been characterized by cryo-EM studies, whereby the binding of haem metabolites to the NTD results in allosteric changes of antibody epitopes,^39^ while linoleic acid binding to the RBD shifts the conformational equilibrium of the S protein to favour down state.^25^ In the case of the S2 site, increases in deuterium exchange observed at the central helical bundle and adjacent regions (peptide 600-613, 946-981, 853-859) may indicate disruption of inter-protomer contacts of the S protein (Figure S4), potentially exacerbated by accommodation of more than one lipid molecule, also supported by our simulations below.

Overall, the deuterium exchange profile of the S protein:lipid A state serves as a reference to differentiate between interactions of the aliphatic lipid tails versus the long polysaccharide chain with the S protein. Comparing the two suggests that deuterium exchange readout from S protein bound states predominantly reports on aliphatic lipid tail interactions with S protein, at three dominant sites spanning the NTD / RBD pockets in S1 and a separate S2 site.

### Analysis of binding of SARS-CoV-2 S1 and S2 to LPS and lipid A

To study the binding between LPS or lipid A with the S protein subunits S1 and S2, we used BN-PAGE followed by western botting, using specific antibodies against His-tag. We have previously shown that binding of *E. coli* LPS to SARS-CoV-2 S protein affects the migration of the protein on BN-PAGE.^19^ In agreement with our previous results, S protein alone migrated at a molecular mass of around 480 kDa (Figure 2A, left panel).^19^ The S1 and S2 subunits migrated at molecular masses of around 242 and 146 kDa, respectively. The S1 and S2 band intensity decreased with increasing doses of LPS which indicates an interaction between the proteins and LPS, as illustrated by the histograms (Figure 2A, right panel). We observed that the change in migration was more marked for S2 when compared with S1. Next, we evaluated the binding of S1 and S2 to lipid A. S protein was included for comparison (Figure 2B). We found that both subunits bind to lipid A, as shown by a decrease of bands at 242 and 146 kDa, for S1 and S2, respectively, and concomitantly a formation of high molecular weight material not entering the gel was observed. Interestingly, lipid A binding to S2 was already significant at 100 μg/ml of lipid A. SDS-PAGE analysis of S, S1 and S2 under denaturing conditions showed homogenous bands (Figure S5A). Using native conditions however, BN-PAGE analysis showed that the preparations of S, S1 and S2 all contained several forms of higher molecular weights when compared with the monomer. This is not unexpected, particularly for the S2 subunit which has been reported to assume a post-fusion trimer structure that is more stable and rigid, both in the absence or the presence of ACE2.^23^ To explore whether the interaction between S2 and LPS was dependent on the pre- and/or post-fusion conformation of S2 we therefore used a mutant-S2 subunit having targeted amino acids substituted with proline residues (F817P, A892P, A899P, A942P, K986P, V987P) in order to prevent fusogenic structural rearrangements in S2.^40^ After incubation with increasing doses of LPS, the proline-substituted S2 protein was analysed on BN-PAGE (Figure S5B) followed by western blot analysis. As above with S2, apart from the monomer at around 146 kDa, the results showed bands of different molecular weights. It was, however, observed that the monomer band at 146 kDa was more defined than seen with the wild type S2 subunit (Figure 2A). Moreover, in the samples with LPS, almost no visible S2 protein was seen in the gel, not even at lower concentrations, suggesting a more pronounced interaction between LPS and S2 in its pre-fusion conformation. In addition, we did not observe any bands at higher molecular weights that were stacked on top of the well, as previously detected for the S2:lipid A mixture (Figure 2B), suggesting the formation of larger aggregates incapable of penetrating the gel during electrophoresis.

**Figure 2:**
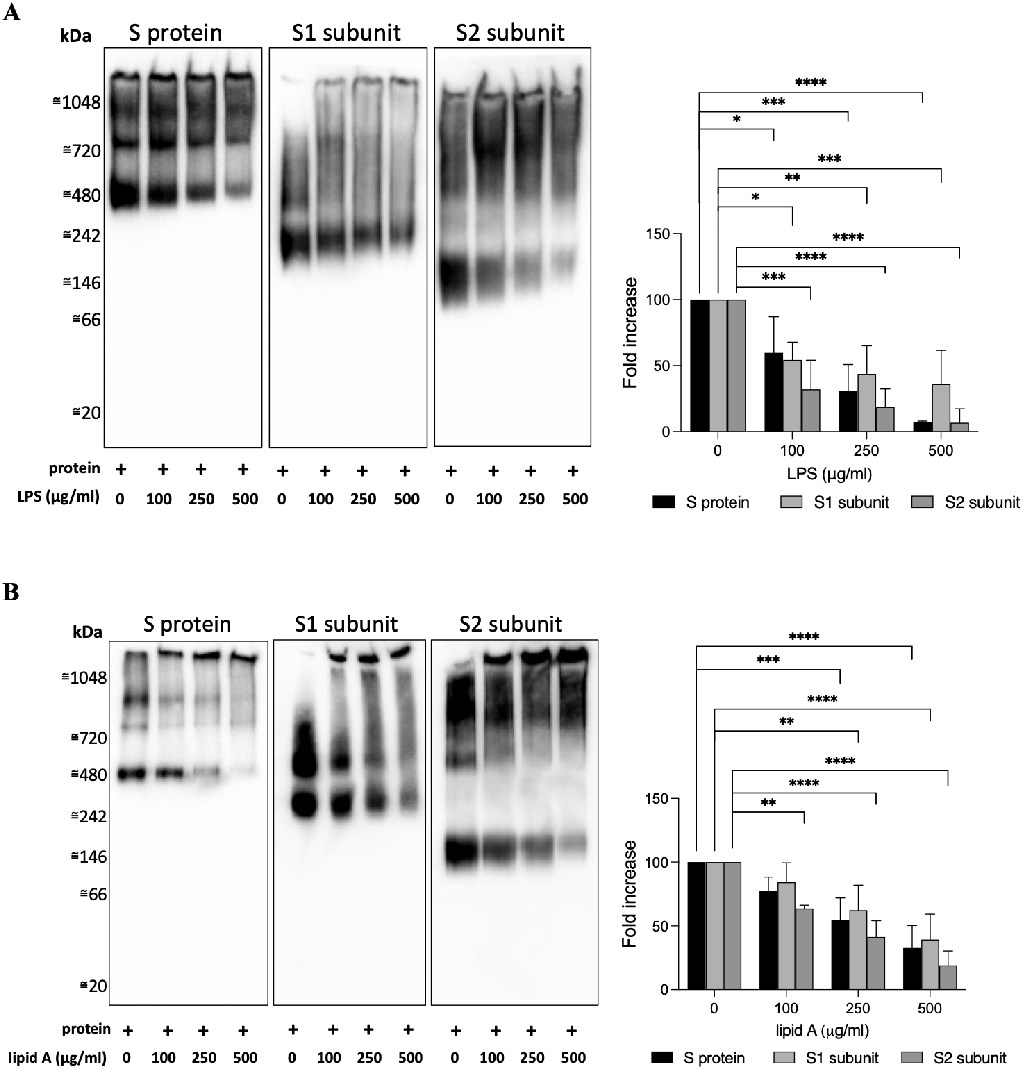
Analysis of binding of S1 and S2 subunits to LPS and lipid. **A**. SARS-CoV-2 S protein or its subunits, S1 and S2, were incubated with (0–500 μg/ml) LPS (A) or lipid A (B), separated using BN-PAGE and detected by western blotting. One representative image of three independent experiments is shown (n=3). The intensity of the bands at 480, 242 and 146 kDa, corresponding to molecular mass of monomeric SARS-CoV-2 S protein or S1 and S2 subunits, respectively, were quantified using Image Lab software 6.1 from Bio-Rad.

### Spike protein binding sites have different affinities for LPS

Our HDXMS data have identified at least three binding sites for LPS and lipid A on the S protein: the NTD, RBD and S2 pockets. This indicates that both S1 and S2 subunits have binding sites for LPS, which was verified experimentally using BN-PAGE. Our previous study showed that LPS binding to the S2 pocket involved residues from both S1 and S2 subunits.^19^ To determine whether LPS binding to this site requires both S1 and S2, we performed MD simulations of LPS bound to this site without the S1 component. MD simulations provide a means to analyse the dynamics of such ligand-protein interactions at atomic resolution. Interestingly, LPS remained stably bound to the isolated S2 domain, as assessed by measurement of the root mean square deviation (RMSD) of the lipid with respect to its initial conformation, which remained low and comparable to the RMSD observed during simulations of the whole S ECD (Figure S6). This suggests that the key residues for LPS binding in this pocket are those found on S2, and residues from S1 may be less crucial for binding. For comparison, simulations of LPS bound to this binding site in a truncated system containing just the S1 subunit revealed LPS detachment from the binding site, and consequently, significantly higher RMSD values. This indicates that LPS binding to the S1 subunit observed in our BN-PAGE experiment does not involve the S2 pocket, but rather involves the pockets in the NTD and RBD only.

Cryo-EM studies revealed that the hydrophobic pockets within the NTD and RBD can accommodate the binding of small hydrophobic molecules, such as polysorbate, linoleic acid and haem metabolites.^25,26,39^ As the lipid A component of LPS contains multiple acyl tails and a large diglucosamine headgroup, how it binds to the NTD and RBD pockets remains unclear. To characterize the dynamics of LPS binding to these domains, we performed unbiased MD simulations whereby a hexa-acylated *E. coli* lipid A molecule was placed nearby the NTD and RBD facing the hydrophobic pockets. We found that in all simulations, the lipid tails spontaneously inserted into the binding sites. The hydrophobic pockets expanded to accommodate the lipid tails, especially on the RBD in which the pocket volume became significantly larger in simulations upon complexation with lipid A (Figure S7). We calculated the solvent accessible surface area (SASA) of individual lipid tails to determine how many of them can bind to the pocket simultaneously. The NTD pocket can accommodate up to five tails, whereas the flexible RBD site can accommodate as many as all six lipid tails of lipid A (Figure 3A-3B) upon pocket expansion. Congruently, we found that the binding of lipid A to RBD results in a lower percentage of its lipid tail atoms being exposed to solvent compared to binding to NTD, suggesting the potential for stronger binding (Figure 3C and 3D). Structural clustering analysis suggests that NTD can accommodate a limited length of each of the lipid tails (approximately 5 carbon atoms distal from the glucosamine units), whereas the RBD can accommodate the entirety of the tails (Figure 3E-3F).

**Figure 3:**
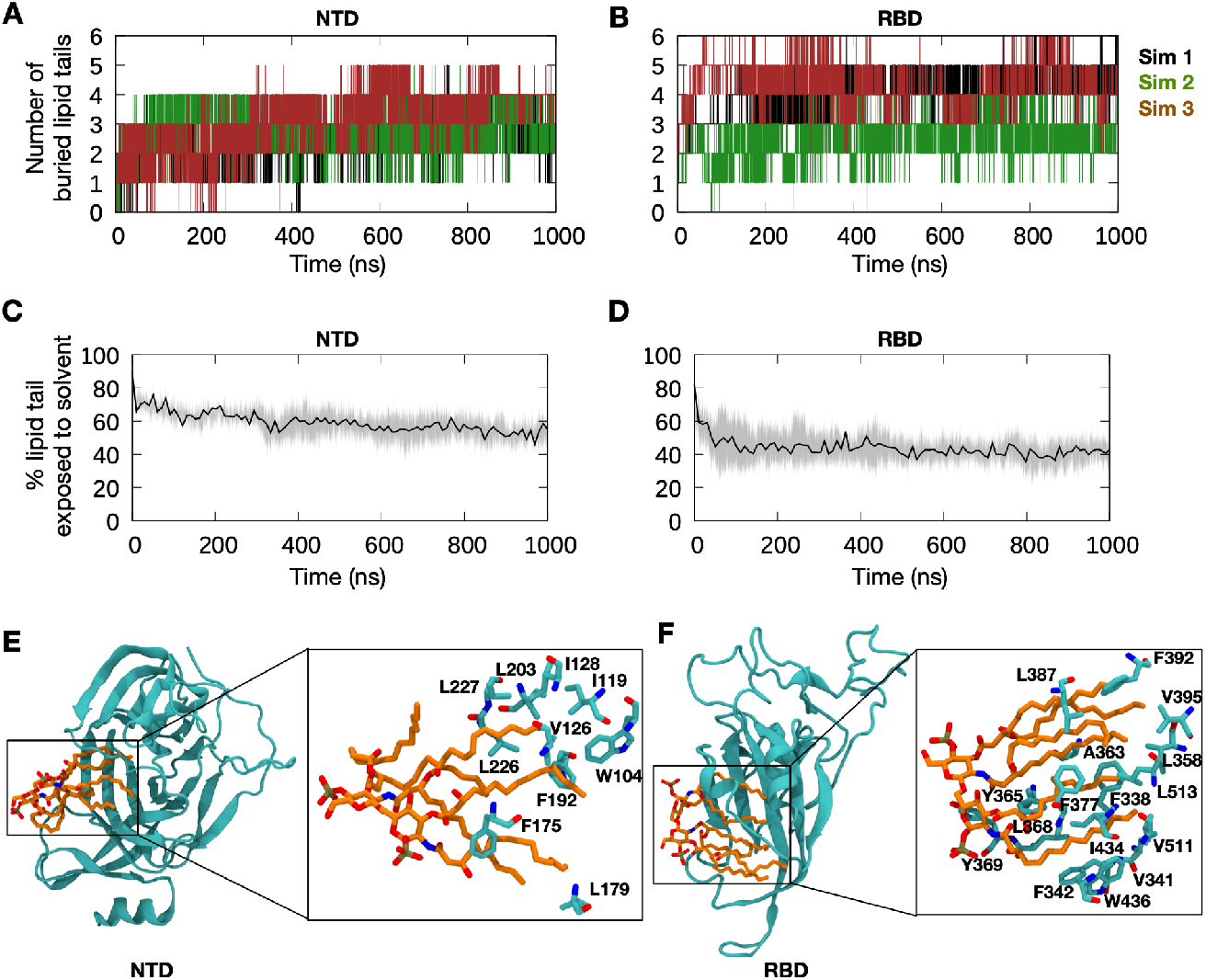
Spontaneous binding of lipid A to S protein NTD and RBD pockets. (A and B) One lipid A molecule was placed nearby the polysorbate binding site on the NTD^26^ or the linoleic acid binding site on the RBD^25^, respectively, and three independent replicas of 1000 ns MD simulations were performed for each system. The number of lipid tails buried within these binding sites were then calculated based on the solvent accessible surface area (SASA) of individual lipid tails. A cut-off of 0.5 nm^2^ was used to categorise the lipid tail as buried. (C and D) Percentage of lipid tail atoms exposed to solvent. Solvent exposed atoms are defined as those found within 0.4 nm of any water molecule. Thick lines show average over 3 independent simulations and shaded areas indicate standard deviations. (E and F) Representative bound states, selected based on cluster analysis. Cluster analysis was performed using 30,000 snapshots generated during the simulations and an RMSD cut-off of 0.4 nm. The central structures of the top clusters are shown. Lipid A is represented in orange and stick representation, while NTD and RBD are represented in cartoon representation in cyan. Enlarged images show hydrophobic residues on the NTD and RBD that interacted with the lipid tails.

While our unbiased MD simulations further validate LPS binding to these pockets on the S1 and S2 subunits, the binding affinity to each pocket remains unknown. To estimate lipid A affinities to all three identified binding sites on the S protein, we performed potential of mean force (PMF) calculations. Steered MD simulations were run to extract lipid A from each site and into bulk solvent, from which snapshots were selected as windows for umbrella sampling (details in Methods). Adequate sampling along the reaction coordinate was achieved, as indicated by the excellent histogram overlap and convergence of each PMF profile (Figure S8). The PMFs suggest that lipid A binds most strongly to the RBD pocket, with a free energy estimated to be around 125 kJ mol^-1^ (Figure 4A). Intriguingly, this is comparable to the value of 135 ± 5 kJ mol^-1^ previously calculated for CD14^29^, which serves as a conduit for LPS in the TLR4 pathway. This result is thus consistent with our previous microscale thermophoresis experiment showing similar *K*_D_ values for LPS interaction with S protein and CD14.^19^ The NTD pocket exhibited a slightly lower binding affinity with an estimated free energy of approximately 95 kJ mol^-1^ (Figure 4B). This agrees with our unbiased simulations showing that lipid A binding to NTD involves a reduced portion of the lipid tails, compared to the RBD which can accommodate the full length of the tails and as many as all six tails simultaneously (Figure 3E and 3F).

**Figure 4:**
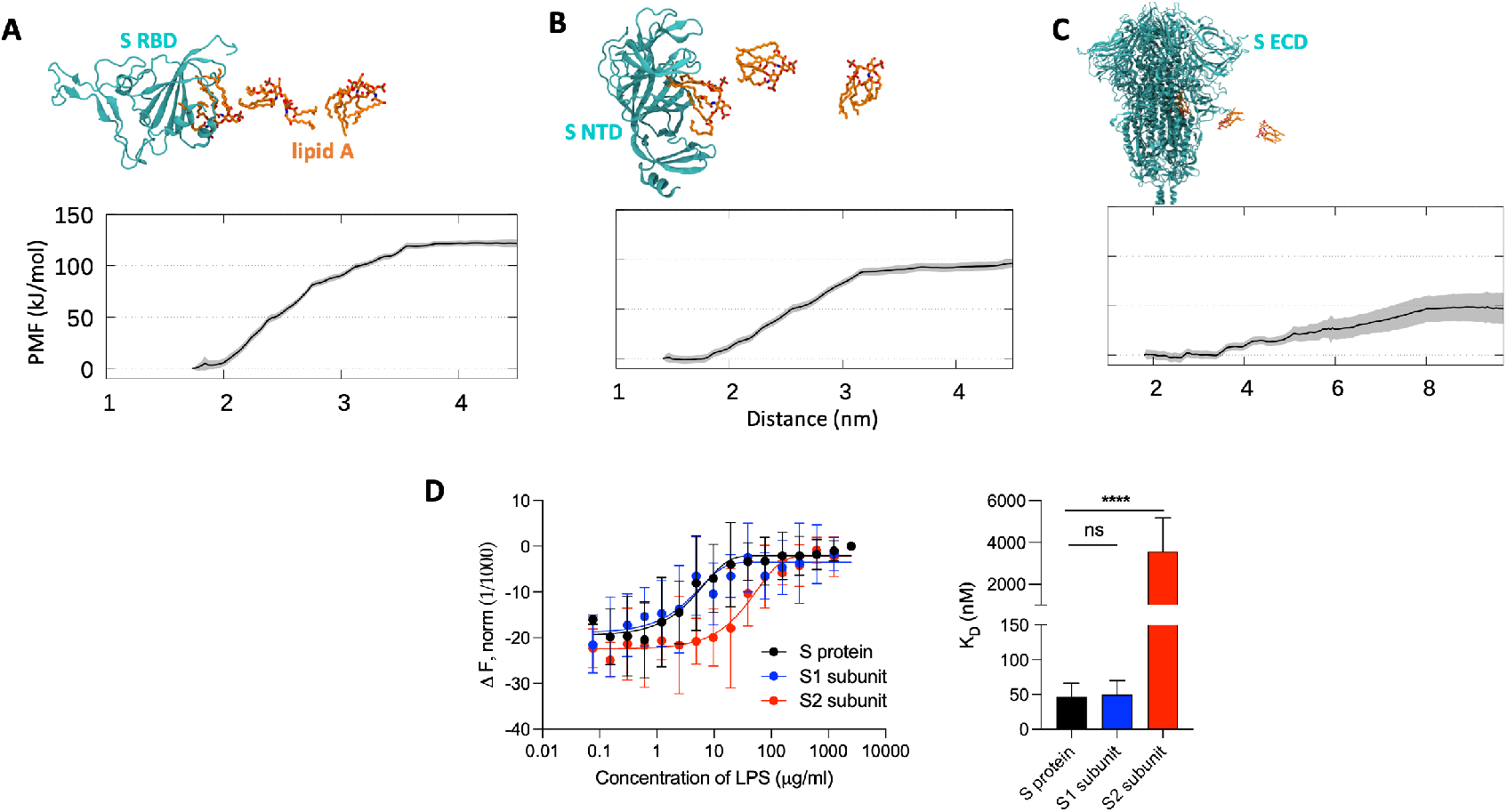
Quantifying lipid A binding to S protein. (A) The calculated potential of mean force (PMF) for lipid A (un)binding to the RBD pocket is shown (bottom panel), with snapshots along the pathway indicated (top panel), based on umbrella sampling simulations. Protein is shown in cyan cartoon representation, while lipid A is shown in orange stick representation. The reaction coordinate is the distance between the centers of mass of lipid A and the protein. Shaded areas indicate standard deviations estimated using bootstrapping. (B and C) Similar PMF calculations for lipid A at the NTD pocket and the S2 pocket, respectively. (D) The binding affinity of LPS to S1 and S2 subunits of S protein was quantified by microscale thermophoresis. Representative binding curves are shown on the left, while K_D_ constant obtained from the curves are summarized with the histograms on the right. K_D_ for S protein is 46.7 ± 19.7 nM, for S1 subunit 50.0 ± 20.0 nM and for S2 3.6 ± 1.6 μM. Data are shown as mean ± SD (n=6).

Finally, our PMF calculations for the S2 pocket yielded a free energy value of around 50 kJ mol^-1^ (Figure 4C) which hence represents the weakest lipid A binding pocket on the S protein. Our previous pocket mapping study using benzene probes on the S protein indicated that the S2 pocket is a deep surface indent, rather than a quintessential binding pocket like the ones on NTD and RBD^37^, hinting at a lower protection of the lipid tails from solvent, and hence weaker affinity. We calculated the percentage of lipid tail exposure to water in each of the umbrella sampling windows for all three PMF calculations. Indeed, the S2 pocket displayed a less dramatic increase in solvent exposure during lipid dissociation, with 60-70% lipid tails buried when bound compared to 80% when in bulk solvent, in contrast with the RBD pocket which is associated with a steep increase from less than 40% to 80% upon unbinding (Figure S9). While the binding affinity to the S2 pocket may seem much weaker than the NTD and RBD pockets, the free energy value is still significant, indicating that a free lipid A molecule would favourably bind to this site. However, in the presence of proteins with higher affinities to lipid A, such as CD14, the lipid A molecule bound equilibrium would presumably shift, suggesting that the S2 pocket could play a role as an intermediary in the TLR4 cascade.

To verify the predicted affinities of LPS to the different S protein binding pockets, we performed microscale thermophoresis (MST) and determined the dissociation constant (K_D_) values for LPS binding to the S1 and S2 subunits independently (Figure 4D). The K_D_ obtained for the S1 subunit was in the same range as for the S protein, 50.0 ± 20.0 and 46.7 ± 19.7 nM, respectively. The low K_D_ value for the S1 subunit is in good agreement with our PMF calculations showing the presence of high-affinity binding pockets for lipid A in the NTD and RBD. Conversely, the K_D_ obtained for the S2 subunit was 3.6 ± 1.6 μM. This higher K_D_ value supports the lower free energy of dissociation determined by our PMF calculations, further corroborating a weaker-affinity binding site for LPS on the S2 subunit.

Intriguingly, our BN-PAGE experiment showed that LPS alters the migration of S2 more prominently compared to S1 (Figure 3), particularly for the prefusion stabilized S2 construct (Figure S5B), suggesting a more pronounced interaction with S2. We note that the concentration of LPS used in all our BN-PAGE experiments is well above the K_D_ values determined by MST (10-50 μM). Due to the large size of the inter-protomeric gap forming the S2 pocket and the partially exposed lipid tails of bound LPS, we hypothesized that more than one LPS molecule could occupy the binding site potentially through lipid-lipid interactions, which would explain the marked changes in migration pattern of the S2 subunit in the presence of LPS. To test this hypothesis, we performed an unbiased MD simulation with three additional lipid A molecules placed outside of the groove, whose centers of mass were at least 2 nm from the bound lipid A. We found that within 300 ns, all three lipid A molecules spontaneously entered the S2 pocket and formed a small lipid A aggregate within the binding site with the already bound lipid A molecule (Figure S10). This aggregate remained stably bound to the S2 pocket for the remainder of the 500 ns simulation. This suggests that the S2 pocket could act as a pool for small LPS aggregates, and may therefore be able to serve a role in forward transfer along the TLR4 relay.

### LPS binding to S protein subunits boosts proinflammatory responses

We have previously shown that S protein, when combined with ultralow-threshold levels of LPS, boosts nuclear factor-kappa B (NF-κB) activation in monocytic THP-1 cells and proinflammatory cytokine release in human blood.^19^ Since both S1 and S2 subunits bind to LPS, we next explored whether these S protein subunits could enhance proinflammatory responses as well. Therefore, we first incubated THP-1 cells with increasing concentrations of LPS (0-1 ng/ml) and a constant amount (5 nM) of S1 or S2. After 20 h, the levels of NF-κB were measured. The results reported in Figure 5A (left panel), show that the S2 subunit alone stimulates NF-κB production. This observation prompted us to investigate whether the preparation could be contaminated by LPS, as has been reported for various commercial S protein preparations.^41^ Indeed, we found that the endotoxin levels were 3.5 and 1401.1 fg/μg in S1 and S2, respectively. It is of note that the level of LPS observed in the S2 preparation does not yield any detectable proinflammatory response in THP-1 cells *per se*; 1 and 10 pg/ml LPS yielded no NF-κB activation (n=3, p=0.9982 and 0.9998, respectively). Nevertheless, taking this confounding endogenous LPS contamination into account, the results demonstrate that exogenously added LPS at ultra-low levels indeed induce a boosting of NF-κB. Surprisingly, no boosting effect was observed for the S1 subunit, either alone or in combination with LPS. Nevertheless, the proinflammatory effect of LPS was not even supressed by the addition of the S1 subunit. It is worth noting that no tested condition was cytotoxic, excluding low cell response to stimuli due to death (Figure 5A right panel). To verify that S2 had a higher proinflammatory effect because of LPS contamination, we performed the experiments on THP-1 cells in the presence of polymyxin B, a well-established neutralizer of LPS.^42^ From the result reported in Figure 5B it is seen that the addition of polymyxin B suppresses the NF-κB activation, both due to S2 alone and to S2 mixed with LPS, indicating an LPS-mediated effect by the S2 preparation. As expected, no visible alteration in NF-κB levels was observed in the case of THP-1 cells treated with the S1 subunit alone, or with LPS and/or polymyxin B. The results with the S2 preparation prompted us to investigate the behaviour of the stabilised prefusion construct, i.e. S2 P-mutant. This variant contained 5.7 fg of LPS /μg of protein, which was comparable to the levels observed in S protein and S1 subunit. Correspondingly, the prefusion stabilized S2 construct showed less intrinsic proinflammatory activity *per se*, but retained a significant ability to boost the responses to ultra-low levels of LPS (Figure S11). Hence, the stabilized S2 protein retained the inflammation boosting activity previously demonstrated for the whole S protein.^19^

**Figure 5:**
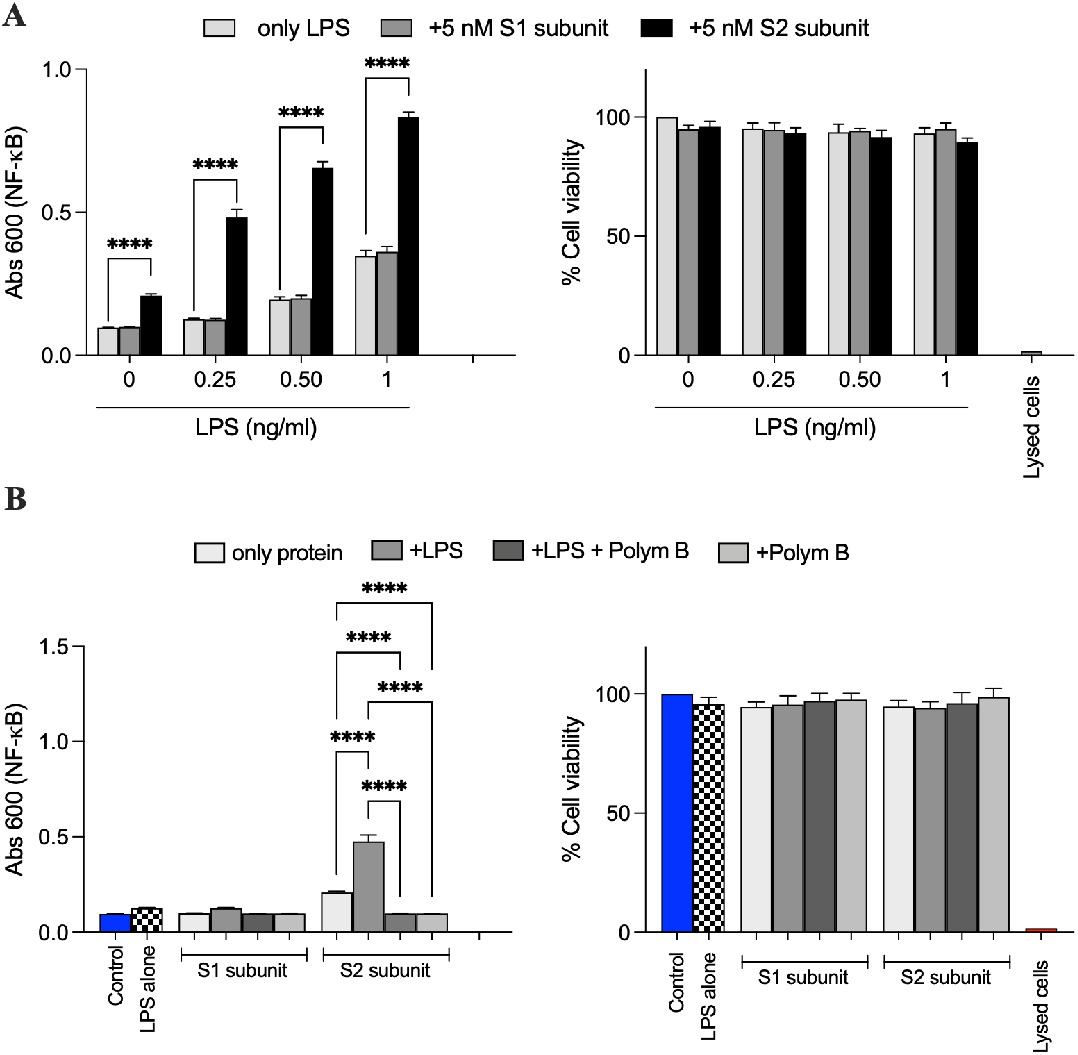
S2 subunit boosts the proinflammatory response to LPS. (A) NF-κB activation (left panel) and cell viability (right panel) were measured in THP-1-XBlue-CD14 cells stimulated with increasing doses (0–1 ng/ml) of LPS and a constant amount (5 nM) of S1 and S2 subunit of S protein. Lysed cells were used as negative control for cell viability. Data are presented as mean ± SD of four independent experiments performed in triplicate (n=4). \*\*\*\**P* ≤ 0.0001, determined using two-way ANOVA with Tukey’s multiple comparisons test. (B) THP-1-XBlue-CD14 cells were treated with 5 nM S1 or S2 subunit, alone or in combination with 0.25 ng/ml LPS and/or 100 μg/ml polymyxin B. After 20 h incubation the NF-κB activation (left panel) and cell viability (right panel) were measured. Lysed cells were used as negative control for cell viability. Data are presented as mean ± SD of four independent experiments all performed in triplicate (n=4). \*\*\*\**P* ≤ 0.0001, determined using ordinary one-way ANOVA with Tukey’s multiple comparisons test.

Using mice reporting NF-κB activation, we previously showed that neither 2 μg LPS nor 5 μg SARS-CoV-2 S protein alone induced a significant proinflammatory response when injected subcutaneously. However, when combined together, a marked boosting of inflammation was observed.^19^ Using a similar setup, we decided to investigate the responses to S1 and S2, and effects on LPS. S1 alone did not yield any measurable NF-κB activation (Figure 6A). S1 when combined with 2 μg LPS resulted in a significant proinflammatory response. S2 alone yielded a proinflammatory response *per se*, which was not unexpected given the detected LPS content in this preparation (Figure S12). Like the *in vitro* results above (Figure 5), addition of polymyxin B to the S2 preparation reduced the proinflammatory response, indicating that LPS was causing the NF-κB activation (Figure S12). Analogously to the *in vitro* data, the prefusion stabilized S2 form showed a significant boosting effect on LPS responses (Figure 6B). Taken together, the results showed that S1 has a boosting effect on LPS-induced inflammation. Indirectly, the results also showed that S2’s inherent proinflammatory effect is due to minute LPS-contaminants, resulting in a boosted NF-κB response to the combination. The proline-stabilized S2 construct, which contained less contaminating LPS, exhibited a low pro-inflammatory activity alone, but significantly boosted the response to 2 μg LPS.

**Figure 6:**
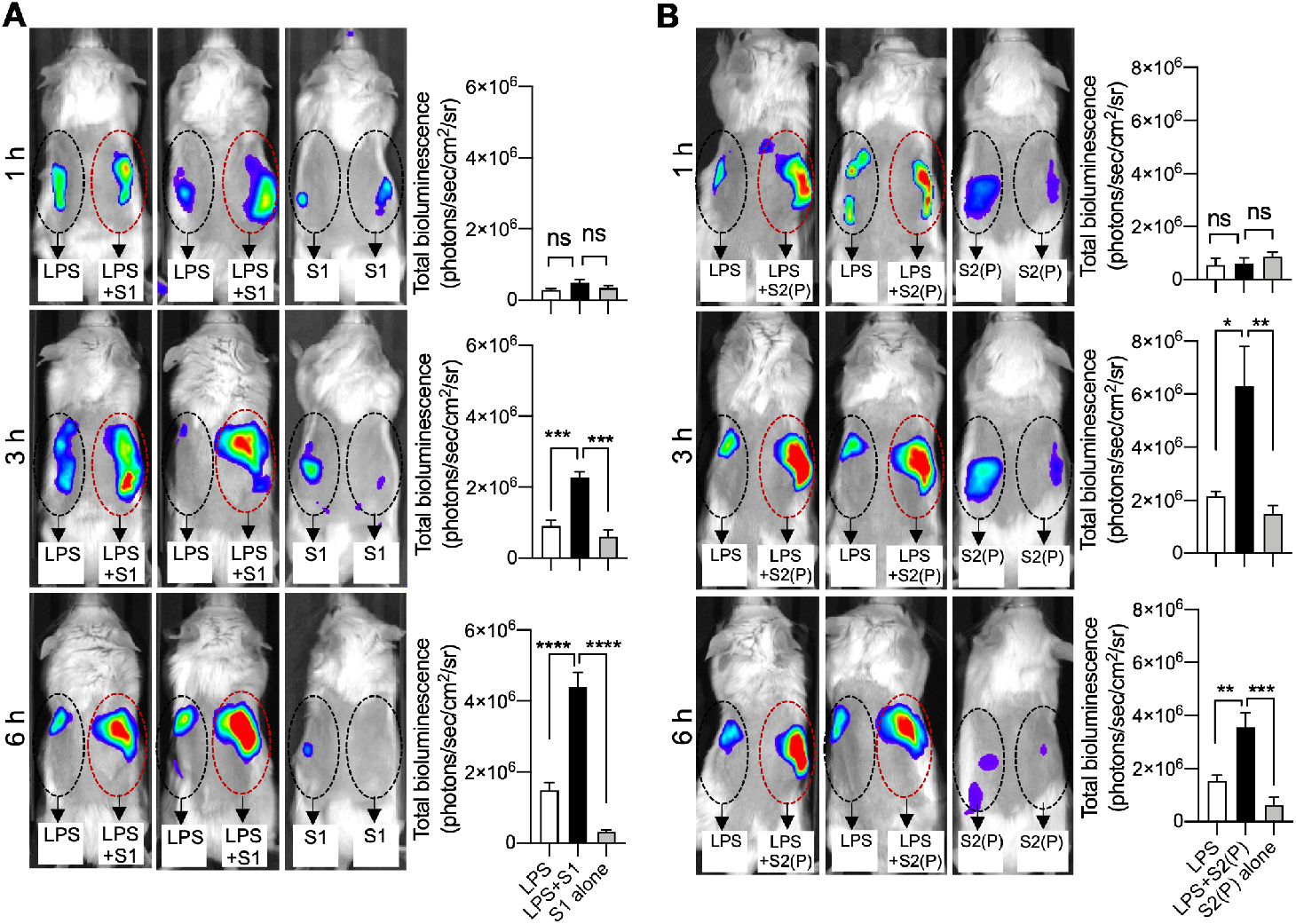
LPS binding to SARS-CoV-2 S protein subunits boosts *in vivo* proinflammatory responses in NF-κB reporter mice. Longitudinal bioimaging of inflammation in NF-κB reporter mice. LPS alone or in combination with subunits S1 (A) or S2 (B) was subcutaneously deposited on the left and right side on the back of transgenic BALB/c Tg(NF-κB-RE-luc)-Xen reporter mice. *In vivo* imaging of NF-κB reporter gene expression was performed using the IVIS Spectrum system. Representative images show bioluminescence at 1, 3 and 6 h after subcutaneous deposition and bar charts show bioluminescence emitted from these reporter mice. Areas of subcutaneous deposition and region of interest for data analysis are depicted as dotted circles. Data are presented as the mean ± SEM (*n* = 4). *P* values were determined using a one-way ANOVA with Dunnett posttest. \**P* ≤ 0.05; \*\**P* ≤ 0.01; \*\*\**P* ≤ 0.001; \*\*\*\**P* ≤ 0.0001; ns, not significant.

It is intriguing that S1 displays a boosting effect on LPS-induced inflammation *in vivo* but not *in vitro*, suggesting additional mechanisms in the former. We hypothesize that one potential reason for this *in vitro*-*in vivo* difference is the interactions with proteases secreted in the tissue during inflammation, such as the neutrophil elastase (NE), that could modify the structure of the S1 subunit and alter its affinity to LPS. To test this hypothesis, we incubated THP-1 cells with the digested and undigested S1 subunit in the absence and presence of LPS. Indeed, S1 mixed with LPS and then digested with NE for 15 minutes was able to increase NF- κB production compared to the mixtures without NE and pre-digested S1 (Figure S13). This supports our hypothesis that LPS binding to the high affinity sites in S1 *in vivo* is compromised by proteolysis during inflammation resulting in an onward transfer to LPS receptors. The proteolytic sites of these enzymes and how they weaken LPS binding, however, would require further studies.

## Discussion

There is growing evidence that bacterial LPS plays a central role in severe COVID-19 complications^15–19,43,44^. Overstimulation of the TLR4 pathway by LPS triggers a hyperinflammatory state that can lead to sepsis and ARDS. In this study, we showed that LPS binds to SARS-CoV-2 S protein at various sites resulting in amplified proinflammatory responses. In the TLR4 pathway, LPS is transferred from the outer membrane of Gram-negative bacteria through a series of receptor proteins to its terminal receptor site, the TLR4:MD-2 complex, which initiates downstream signal activation^45–47^. An affinity gradient exists between LPS and these receptors to favour a one-way transfer that culminates at TLR4:MD-2^29^. Here, we propose how the S protein fits into the LPS transfer cascade (Figure 7). S protein disaggregates LPS micelles^19,28^ via high affinity binding to the S1 pockets on the NTD and RBD (“high-affinity, low-capacity” binding sites), which are positioned at the outermost part of the S protein and the virus. The NTD and RBD pockets have similar LPS binding affinities to the CD14 receptor,^29^ which *in vitro* results in unfavourable onward transfer. However, the presence of proteases secreted during inflammation *in vivo* could compromise LPS binding resulting in LPS dissociation and transfer to other receptors. Additionally, the S1 pockets can also bind to various endogenous hydrophobic molecules such as lipids and metabolites,^25–27^ which could displace LPS and promote subsequent transfer events. Conversely, the S2 site has a weaker binding affinity and is considerably larger than the S1 pockets, which allows for simultaneous binding of multiple LPS molecules (low-affinity, high-capacity” binding site). Thus, the S2 pocket could act as a “sink” for LPS upon disaggregation, and the lower affinity promotes onward transfer to CD14 along the TLR4 relay. Collectively, the S protein enhances the LPS transfer cascade by acting as an additional delivery mediator to the LPS receptors.

**Figure 7:**
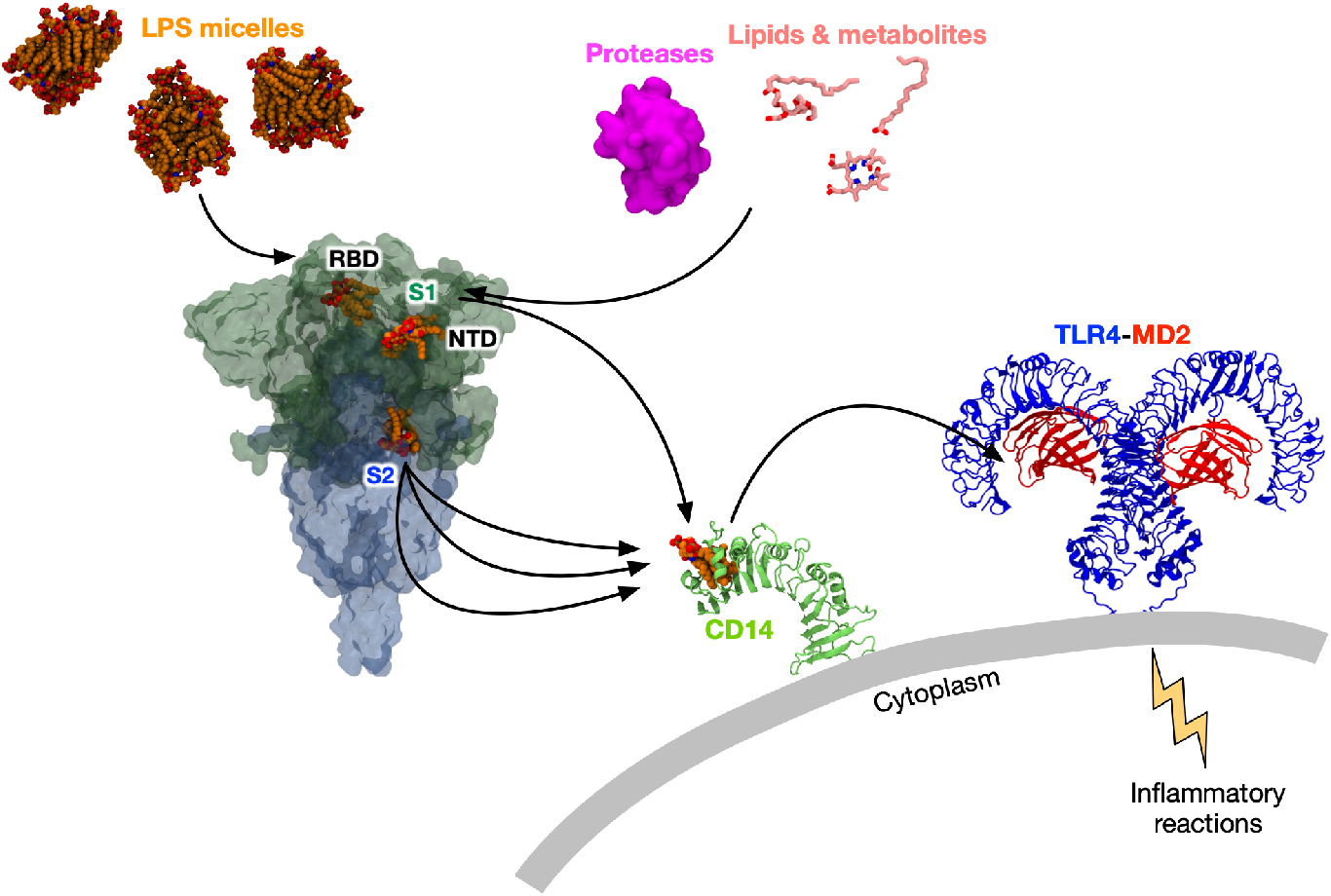
Proposed model for S protein boosting effect on LPS-mediated proinflammatory response. In the presence of S protein, LPS micelles disaggregate due to binding to high-affinity, low-capacity sites on the S1 subunit. These binding pockets have similar affinities to CD14, resulting in unfavourable onward transfer. However, the presence of proteases *in vivo* that may compromise LPS binding, as well as endogenous hydrophobic molecules such as lipids and metabolites that may displace bound LPS, result in transfer to CD14. Free LPS molecules can also bind to the S2 subunit, whose low-affinity, high-capacity binding pocket can accommodate more than one LPS molecule simultaneously. The lower binding affinity would also facilitate downstream LPS transfer to CD14. Subsequently, LPS is transferred to the TLR4:MD-2 complex, which triggers proinflammatory reactions. Thus, the S protein acts as an intermediate in the LPS receptor transfer cascade.

Nevertheless, it is possible that the S protein itself induces hyperinflammation and viral sepsis independent of bacterial LPS. Previous studies suggested that SARS-CoV-2 S protein binds TLR4 directly and activates related immune responses.^48,49^ Our experiment with polymyxin B, an LPS-depleting agent, however, suggests that those results could be caused by LPS contamination. Indeed, in the presence of polymyxin B, the inflammatory boosting effect by the S2 subunit of the S protein observed in THP-1 cells and NF-κB reporter mice was eliminated, indicating that LPS is categorically essential, and that the S protein alone is not able to activate the TLR4 pathway. Moreover, our findings that the intrinsic proinflammatory activity of the two different S2 preparations is correlated to presence of minute LPS contaminants elegantly confirmed this assumption. Indeed, a recent study showed that commercially produced S protein preparations contain varying concentrations of LPS, which correlate with their abilities to induce cytokine expression in blood cells.^41^ Similarly to our results, the cytokine production by these S protein reagents was abrogated by polymyxin B. This further corroborates our proposed mechanism whereby the S protein is an intermediary rather than a direct causality of hyperinflammation. It is worth noting that the S protein is not the only SARS-CoV-2 protein associated with inflammatory responses; a recent study revealed that the envelope (E) protein is recognised by the TLR2 receptor and induces the production of inflammatory cytokines.^50^

To date, around 2.7 million sequences of SARS-CoV-2 genome have been deposited on the Global Initiative on Sharing All Influenza Data (GISAID) platform.^51^ As SARS-CoV-2 is a rapidly mutating RNA virus, almost all of the residues on the S protein have been detected to mutate at least once since its discovery in January 2020.^52^ We calculated the percentage of occurrence of amino acid changes from hydrophobic to non-hydrophobic residues for the LPS binding residues and found a value of well below 0.01% for all positions in the three binding pockets (Figure 8A). This indicates that the LPS binding residues are highly conserved in emerging SARS-CoV-2 variants and are unlikely to mutate in the future. Hence, the LPS binding pockets could be attractive targets for development of therapeutics. Interestingly, when compared to sequences from other related beta-coronaviruses, most of these residues are also conserved across different species (Figure 8B), suggesting a potentially universal LPS binding capability by these coronaviruses S proteins. We previously showed that the S protein from SARS-CoV interacts with LPS similarly to the S protein from SARS-CoV-2;^19^ further studies would be required to determine LPS interactions with S proteins from other coronaviruses.

**Figure 8:**
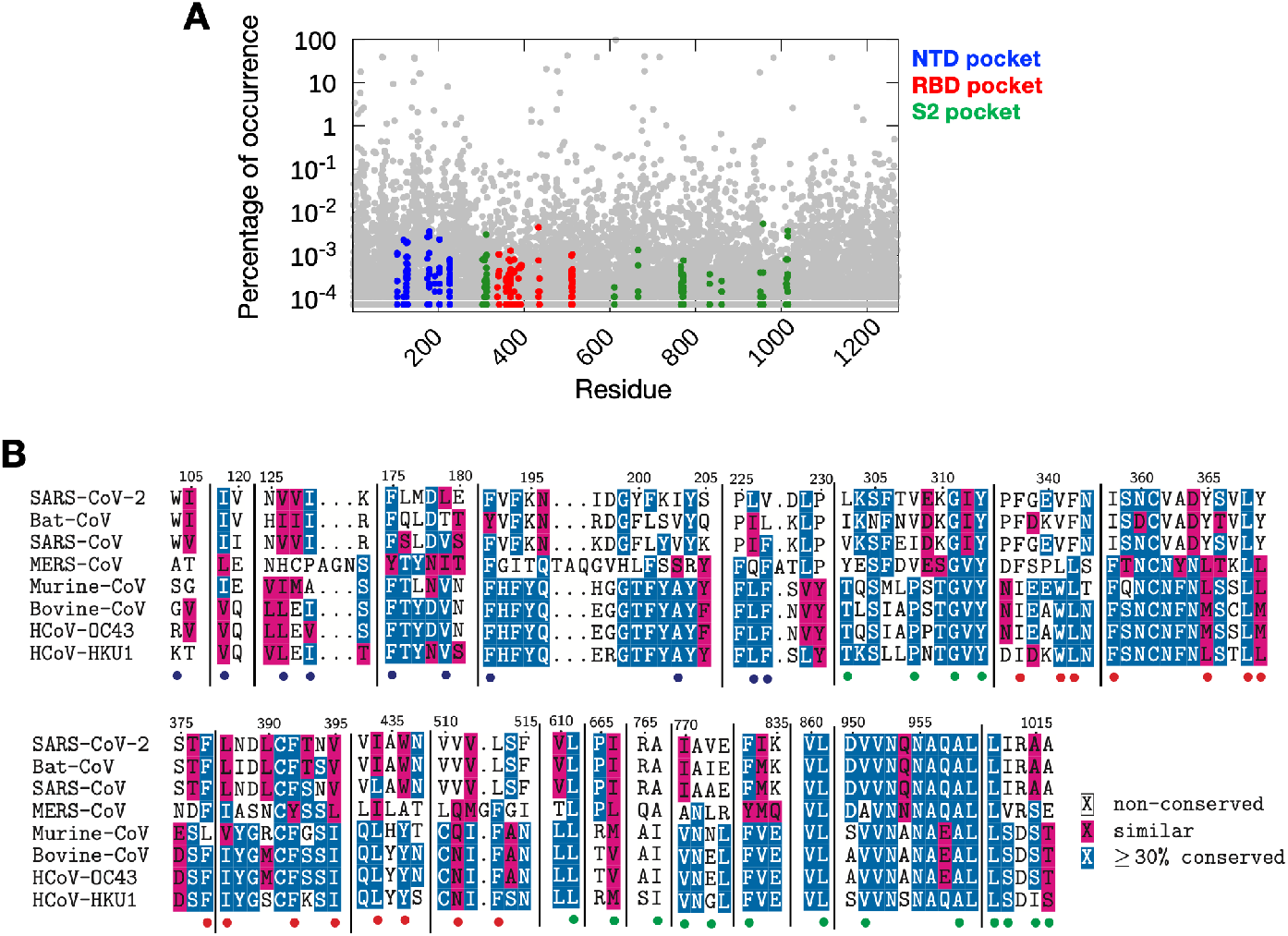
Sequence conservation of LPS binding residues. (A) The percentage of occurrence of amino acid changes at each position of the S protein based on sequences of SARS-CoV-2 deposited in the GISAID initiative database.^3^ The percentages for all mutations are shown in grey, while the percentages for mutations on residues predicted to be involved in LPS binding to non-hydrophobic residues are shown in blue (NTD pocket), red (RBD pocket) and green (S2 pocket). The latter occur at well below 0.01% in all reported sequences to date. Data was taken in August 2021 when 2,701,750 sequences had been reported. (B) Multiple sequence alignment of SARS-CoV-2 S protein with S proteins from other beta-coronaviruses showing conservation of predicted LPS binding residues. Residues that are more than 30% conserved are shaded in blue, while similar residues are shaded in red. LPS binding residues from the NTD, RBD and S2 pockets are highlighted by the blue, red and green circles, respectively. S proteins from different species of coronaviruses with at least 30% sequence identity to SARS-CoV-2 S were selected. UniProt IDs: SARS-CoV-2, P0DTC2; Bat-CoV, Q3LZX1; SARS-CoV, P59594; MERS-CoV, K9N5Q8; Murine-CoV, P11224; Bovine-CoV, P25191; HCoV-OC43, P36334; HCoV-HKU1, Q5MQD0.

The binding of LPS to the NTD and RBD is interesting in itself due to the instrumental role of these subdomains in immune evasion of the SARS-CoV-2. The binding of haem metabolites to the NTD pocket has been shown to yield profound conformational changes through an allosteric mechanism that inhibits neutralizing antibodies targeting the NTD^39^. As LPS binds to the same pocket, it is possible that it could also alter the conformation of the antigenic supersite found on the NTD^53^. In the RBD pocket, linoleic acid binding bridges two adjacent RBDs^25^, hence favouring the RBD down state. Similarly, the charged headgroup of LPS protrudes out of the binding cavity and could potentially interact with a neighbouring RBD and shift the open-closed equilibrium of the S protein. The RBD is better shielded by glycans in its down conformation^54^; therefore, a shift towards the down state by LPS binding could reduce accessibility to antibody epitopes and negatively impact neutralization. Various mutations on the NTD and RBD found in SARS-CoV-2 variants promote accelerated community transmission and vaccine breakthrough^55,56^. As described above, none of the mutations recorded to date appear in any of the LPS binding pockets, implying that the S proteins from these variants are competent to bind LPS. These mutations, along with the effect of LPS binding on antibody epitopes, could therefore synergistically contribute towards immune evasion.

The evolutionary origin of LPS binding capacity by the SARS-CoV-2 S protein remains an enigma. From a structural standpoint, the presence of partially exposed hydrophobic patches that allow for LPS binding is counterintuitive. It is worth noting that some of the residues in the S2 pocket that we have shown interact with LPS acyl tails form the central core of the S protein post-fusion structure^23^. That these residues become more exposed to solvent upon S1 shedding suggests that they may be a part of the “spring-loaded” S2 core that drives the pre-to post-fusion conformational transition. The LPS binding site could therefore arise as a side effect of the inclusion of these metastable features on the S protein structure required for its function during viral infection.

The LPS binding sites could also have emerged to promote direct bacteria-virus interaction. Interkingdom synergy between bacteria and viruses at the host-pathogen interface plays monumental roles in various co-infections^57^. Direct interactions between enteric viruses and bacteria through attachment to surface LPS improves viral host cell adherence and environmental stability^58^, while promoting genetic recombination between viruses to restore viral fitness^59^. Similarly, influenza virus interacts directly with the bacterial surface and increases bacterial adherence to respiratory cells, worsening pathogenesis^60^. It is thus possible that SARS-CoV-2 has evolved to bind Gram-negative bacteria via S protein-LPS interactions to modulate certain aspects of its viral life cycle; however, the exact role of this interaction is likely to be complex and would require further studies.

In conclusion, we report in detail the molecular mechanism of interaction between the SARS-CoV-2 S protein and bacterial LPS, suggestive of how this interaction leads to enhanced TLR4 mediated inflammatory responses. Our study also provides novel therapeutic avenues for drug development against hyperinflammation observed in severe COVID-19 cases.

## Acknowledgement

This work was supported by BII (A*STAR) core funds. P.J.B. and F.S. gratefully acknowledge support from grant FY21_CF_HTPO SEED_ID_BII_C211418001 funded by A*STAR. Simulations were performed using computational resources of the petascale computer cluster ASPIRE-1 at the National Supercomputing Centre of Singapore (NSCC), the A*STAR Computational Resource Centre (A*CRC), and the supercomputer Fugaku provided by RIKEN through the HPCI System Research Project (Project ID: hp200303). A.S., J.P., and G.P. acknowledge support by grants from the Swedish Research Council (project 2017-02341, 2020-02016), Edvard Welanders Stiftelse and Finsenstiftelsen (Hudfonden), the Torsten Söderberg, Crafoord, and Österlund Foundations, Stiftelsen Lars Hiertas Minne, the Royal Physiographic Society of Lund, and the Swedish Government Funds for Clinical Research (ALF). The authors thank Ann-Charlotte Strömdahl for excellent technical assistance with THP-1 cells.

## Author contributions

P.J.B and A.S. designed the study. F.S. performed molecular modelling and simulations. P.R. conducted HDXMS experiments. G.P. conducted BN-PAGE and *in vitro* NF-κB cell activation assays. M.P. performed mouse inflammation model and *in vivo* bioimaging. J.P. performed MST binding assay. P.J.B, A.S., and G.S.A. supervised the experiments and simulations. All authors drafted and finalised the manuscript.

## Materials and Methods

### Material

Lipopolysaccharides from *E. coli* O111:B4 (catalog no. L3024) and lipid A from *E*.*coli* F583 (Rd mutant, catalog no. L5399) were purchased from Sigma Aldrich. Deuterium oxide was from Cambridge Isotope Laboratories (Tewksbury, MA). All reagents and chemicals were research grade or higher and obtained from Merck-Sigma-Aldrich (St. Louis, MO). Mass Spectrometry grade acetonitrile, formic acid and water were from Fisher Scientific (Waltham, MA); and trifluoroacetic acid (TFA), sequence analysis grade from Fluka BioChemika (Buchs, Switzerland).

### S protein expression and purification

S protein expression and purification for use in HDXMS experiments was carried out as described in Raghuvamsi et al.^34^ Briefly, a Pfastbac expression vector containing codon optimised SARS-CoV-2 S protein sequence (16-1208; Wuhan-Hu-1) for insect cell expression was used to express S protein in Spodoptera Sf9 cells. The S protein sequence contain two mutated sites with RRAR (682-685) into DDDDK and residues KV (986-987) into PP, along with HRV 3C site, 8x-Histidine tag, and a streptavidin tag at the C-terminus. Sf9 cells transfected by following were used to express the S protein by following bac-to-bac baculovirus expression system (Thermo Fisher, SG). The purified bacmid of SARS-CoV-2 was used for transfection using cellfectin (Thermo Fisher, SG). The viral stocks obtained from transfection were amplified and expressed for protein production. Cell culture was harvested on fourth day after transfection and supernatant was affinity purified using a 5ml HisTrap excel column (Cytiva, SG). The affinity purified sample was subjected to size exclusion chromatography using HiLoad 16/60 superdex 200 pre-equilibrated with 20mM Hepes, pH 7.5, 300mM NaCl, 5% glycerol. Peak fractions from SEC separated sample was concentrated using a 100 kDa cut-off concentrator (Sartorius, GE) with PBS buffer (pH 7.4). Purity of protein samples was confirmed by SDS-PAGE.

SARS-CoV-2 S, SARS-CoV-2 S1 and two variants of SARS-CoV-2 S2 proteins, produced by ACROBiosystems (USA), were used for BN-PAGE, MST and to stimulate THP-1 cells and for *in vivo* experiments. The sequence of SARS-CoV-2 S protein contains AA Val 16 - Pro 1213 (Accession # QHD43416.1 (R683A, R685A)), the sequence of SARS-CoV-2 S1 protein contains AA Val 16 - Arg 685 (Accession # QHD43416.1), the sequence of native SARS-CoV-2 S2 protein contains AA Ser 686 - Pro 1213 (Accession # QHD43416.1), while its mutant contains beside AA Ser 686 - Pro 1213 also proline substitutions (F817P, A892P, A899P, A942P, K986P, V987P) (Accession # QHD43416.1). All proteins were expressed in human 293 cells (HEK293) and purified exploiting polyhistidine tag at the C-terminus. The proteins were lyophilized from a 0.22 μm filtered solution in 50 mM Tris, 150 mM NaCl, pH 7.5. Lyophilized products were reconstituted in endotoxin free water, aliquoted and stored at – 80 °C according to the manufacturer’s protocol. The purity was >85 for full length SARS-CoV- 2 S, >90% for SARS-CoV-2 S1 protein and >95% SARS-CoV-2 S2 protein.

### Deuterium labelling and quench conditions

Deuterium labelling was performed on free S protein solubilized in PBS at pH7.4 incubated at 37°C in PBS buffer reconstituted in 99% D_2_O attaining at final concentration of 90%. Similarly, S protein incubated with lipid A or LPS molecules dissolved in PBS (pH 7.4, LA 0.5% DMSO) at a molar ratio of 1:10 for 30 minutes (S protein monomer:lipid A/LPS assuming molecular weight of 4 kDa for lipid A and 10 kDa for LPS, respectively) were subjected to deuterium labelling. All the deuterium labelling reactions were performed for 1, 10, or 100 min before quenching the reaction with a prechilled quench buffer. Quench buffer containing 1.5 M GnHCl and 0.25 M Tris(2-carboxyethyl) phosphine-hydrochloride (TCEP-HCl) was added to lower the pH to 2.5. Upon addition on quench buffer the reaction mixture was incubated on ice for 1 min followed by online pepsin digestion.

### Mass Spectrometry and peptide identification

∼100 pmol of quenched reaction mixture was injected onto the nanoUPLC HDX sample manager (Waters, Milford, MA). Online pepsin digestion was performed on the injected samples using immobilized Waters Enzymate BEH pepsin column (2.1 × 30 mm) at a flow rate of 100 μl/min in 0.1% formic acid in water. Simultaneously, the pepsin proteolyzed peptides were trapped in a 2.1 × 5 mm C18 trap (ACQUITY BEH C18 VanGuard Pre-column, 1.7 μm, Waters, Milford, MA). Following pepsin digestion elution of proteolyzed peptide was carried out using acetonitrile gradient of 8 to 40 % in 0.1 % formic acid at a flow rate of 40 μl min^-1^ into reverse phase column (ACQUITY UPLC BEH C18 Column, 1.0 × 100 mm, 1.7 μm, Waters) pumped by nanoACQUITY Binary Solvent Manager (Waters, Milford, MA). Resolved peptides were ionized using electrospray ionization mode and sprayed onto SYNAPT G2-Si mass spectrometer (Waters, Milford, MA). Detection and data acquisition of eluted and resolved peptides was done in HDMS^E^ mode with ion mobility settings set to 600 m/s wave velocity and 197m/s transfer wave velocity. Mass spectra within a range of 50-2000 Da were acquired for 10 minutes with mass spectrometer in positive ion mode. A constant 25 V cone voltage was applied and for trap and transfer low collision energies of 4 V and 2 V was used. High collision energy was ramped from 20-45 V. For lockspray correction, 200 fmol μl^−1^ of [Glu^1^]-fibrinopeptide B ([Glu^1^]-Fib) was injected to at a flow rate of 5 μl/min.

Peptide sequences were identified using Protein Lynx Global Server v3.0 from the mass spectra of undeuterated protein samples. Peptide sequence search was carried out on a separate sequence database of S protein along with purification tag. No specific protease and variable N-linked glycosylation modification were chosen as search parameters for sequence identification. Further peptides were filtered using minimum intensity cutoff of 2500 for product and precursor ions, minimum products per amino acids of 0.2 and a precursor ion mass tolerance of <10 ppm using DynamX v.3.0 (Waters, Milford, MA). Peptides present in at least two out of three undeuterated samples were used for HDXMS. All deuterium exchange reactions were performed in triplicates and no back-exchange corrections were carried out for reported deuterium uptake values. A significance cutoff of ± 0.5 Da was determined from the standard deviation of the average deuterium exchange values from technical replicates and most peptides showed a standard deviation within ± 0.3 Da.

### SDS-PAGE

The purity of SARS-CoV-2 S, S1, S2 and S2 P-mutant subunits was evaluated by SDS-PAGE (Supplementary Fig. S3). Briefly 2 μg of each protein was diluted in loading buffer and loaded on 10–20% Novex Tricine pre-cast gel (Invitrogen, USA). The run was performed at 120 V for 1 h. The gel was stained by using Coomassie Brilliant blue (Invitrogen, USA). The image was obtained using a Gel Doc Imager (Bio-Rad Laboratories, USA).

### Blue Native (BN)-PAGE

Two μg SARS-CoV-2 S1 or S2 subunit was incubated with 0.1, 0.25 or 0.5 mg/ml of *E. coli* LPS or lipid A for 30 min at 37 °C in 20 μl as final volume. At the end of the incubation the samples were separated under native conditions on BN-PAGE (Native PAGE BisTris Gels System 4–16%, Invitrogen) according to the manufacturer’s instructions. Then, the material was transferred to a PVDF membrane using the Trans-Blot Turbo system (Bio-Rad, USA). Primary antibodies against the His-tag (1:2000, Invitrogen) were followed by secondary HRP conjugated antibodies (1:2000, Dako, Denmark), for detection of both proteins. The bands of the proteins were visualized by incubating the membrane with SuperSignal West Pico Chemiluminescent Substrate (Thermo Scientific, Denmark) for 5 min followed by detection using a ChemiDoc XRS Imager (Bio-Rad). LPS as well as lipid A were purchased from Sigma (USA). In another set of experiments, 2 μg of S2 P-mutant was incubated with increasing doses of *E. coli* LPS (0.1–0.5 mg/ml) for 30 min at 37 °C in 20 μl as final volume. At the end of incubation samples were mixed with loading buffer and analysed as described above. All experiments were performed at least 3 times.

### Microscale thermophoresis

Forty μg SARS-CoV-2 S, S1 or S2 protein were labelled by Monolith NT Protein labelling kit RED – NHS (Nano Temper Technologies, Germany) according to the manufacturer’s protocol. 5 μl of 20 nM labeled proteins were incubated with 5 μl of increasing concentrations of LPS (250–0.007 μM) in 10 mM Tris at pH 7.4. Then, samples were loaded into standard glass capillaries (Monolith NT Capillaries, Nano Temper Technologies) and the MST analysis was performed (settings for the light-emitting diode and infrared laser were 80%) on a NanoTemper Monolith NT.115 apparatus (Nano Temper Technologies, Germany). Results shown are mean values ± SD of six measurements.

### Unbiased molecular dynamics simulations

All-atom MD simulations were performed to elucidate lipid A/LPS binding mechanism to the pockets on S1 and S2 subunits of the S protein. For the S1 pockets, a hexa-acylated *E. coli* lipid A molecule was placed 3 nm away from the S protein NTD (residue 14-305) and RBD (residue 335-525) facing the polysorbate and linoleic acid binding sites,^25,61^ respectively. For the S2 pocket, we performed simulations of *E. coli* LPS bound to the pocket with S1 subunit (residue 14-685) and S2 subunit (686-1162) independently. Additionally, we ran simulations of multiple lipid A binding, whereby three lipid A molecules were placed outside of the inter-protomeric groove near the S1/S2 cleavage site at a center of mass distance of at least 2 nm away from the bound lipid A. The initial coordinates of *E. coli* lipid A and LPS were obtained from CHARMM-GUI LPS modeller,^62^ while the initial coordinates of the S protein were extracted from the cryo-EM structure of S ECD in the closed state (PDB: 6XR8)^23^ with missing loops constructed using Modeller version 9.21.^63^ Protein and lipid were parameterized using the CHARMM36 forcefield.^64^ The systems were solvated with TIP3P water molecules and 0.15 M NaCl salts. Steepest descent minimisation and a short 125 ps equilibration simulation were performed, following the standard CHARMM-GUI protocols.^65^ Three replicates of 1 μs simulations were performed for each of the S1 pockets, while three replicates of 500 ns simulations were performed for the S2 pocket, with different initial velocities. Coupling to a Nosé-Hoover thermostat^66,67^ was used to maintain a temperature of 310 K, whilst isotropic coupling to a Parrinello-Rahman barostat^68^ was used to maintain a 1 atm pressure. Coulombic interactions were computed using the smooth particle mesh Ewald (PME) method^69^, and the van der Waals interactions were truncated at 1.2 nm with a force smoothing function applied between 1.0 to 1.2 nm. A 2-fs time step was used with constraints applied on all covalent bonds involving hydrogen atoms using the LINCS algorithm.^70^ All simulations were performed using GROMACS 2018^71^ and the trajectories visualised in VMD.^72^ Most analyses were performed using GROMACS tools, while the pocket volume was calculated using MDPocket.^73^

### PMF calculations

To estimate the binding affinity of lipid A to each binding site on the S protein, PMF calculations were performed along a reaction coordinate parallel to the lipid dissociation pathway from the binding pockets. For the S2 binding site, the docked lipid A complex ^19^ was used as starting coordinates. For the NTD and RBD binding sites, cluster analysis was performed on the unbiased simulations described above, and the central structures of the top clusters were used. Steered MD simulations were first performed to generate the trajectories of lipid dissociation. Lipid A was pulled away from the binding pockets along an axis perpendicular to the protein surface at a constant velocity of 0.1 nm ns^-1^ using an elastic spring with a force constant of 1,000 kJ mol^-1^ nm^2^ applied to its centre of mass. Positional restraints with a force constant of 1,000 kJ mol^-1^ nm^2^ were applied to the backbone atoms of the protein throughout. From these trajectories, 50 umbrella sampling windows were selected based on the distance between the centers of mass of the lipid and protein along the reaction coordinate with a separation of 0.1 nm between windows. Subsequently, a 100 ns simulation was performed for each window with the centre of mass of lipid A restrained in the vector of the reaction coordinate with a force constant of 1,000 kJ mol^-1^ nm^2^. No restraint was applied to the protein. The GROMACS *gmx wham* tool based on the weighted histogram analysis method (WHAM)^74^ was used to compute the PMF from the umbrella sampling simulations. Histogram overlaps were plotted to determine any poor sampling along the reaction coordinate, for which additional simulations were performed. 100 bootstrap trials were performed for each PMF calculation to estimate the statistical errors. Convergence analysis was performed by generating PMF profiles using increasing amounts of simulation sampling time.

### NF-κB activation assay

THP1-XBlue-CD14 reporter cells (InvivoGen, San Diego, USA) were seeded in 96 well plates (180,000 cells/well) in phenol red RPMI, supplemented with 10% (v/v) heat-inactivated FBS and 1% (v/v) Antibiotic-Antimycotic solution. Cells were treated with 5 nM SARS-CoV-2 S1, S2 or S2 P-mutant subunits in the presence of increasing doses (0.25-1 ng/ml) of LPS (Sigma, USA). Then, the cells were incubated at 37 °C for 20 h. At the end of incubation, the NF-κB activation was analysed according to the manufacturer’s instructions (InvivoGen, San Diego, USA), i.e. by mixing 20 μl of supernatant with 180 μl of SEAP detection reagent (Quanti-BlueTM, InvivoGen), followed by absorbance measurement at 600 nm. In another set of experiments, cells were treated with 5 nM SARS-CoV-2 S1 and S2 protein in the presence of 0.25 ng/ml LPS and 100 μg/ml polymyxin B. The experiment was performed as described above. Data shown are mean values ± SD obtained from at least four independent experiments all performed in triplicate.

### MTT assay

The viability of cells exposed to different treatments was evaluated by adding 0.5 mM Thiazolyl Blue Tetrazolium Bromide (MTT) to the cells remaining from NF-κB activation assay and incubating the cells at 37 °C for 2 h. Next, cells were centrifuged at 1000 g for 5 min and the medium was removed. The formazan salts were solubilized by adding100 μl of 100 % DMSO (Duchefa Biochemie, Haarlem) to each well. Absorbance was measured at a wavelength of 550 nm. Cell survival was expressed as percentage of viable cells in the presence of different treatment compared with untreated cells. Lysed cells were used as positive control. Data shown are mean values ± SD obtained from at least four independent experiments all performed in triplicate.

### Mouse inflammation model and in vivo imaging

BALB/c tg (NF-B-RE-Luc)-Xen reporter mice (Taconic Biosciences, Albany, NY, USA, 10– 12 weeks old) were used to study the immunomodulatory effects of SARS-CoV-2 S protein subunits alone or in combination with LPS. Using a trimmer, hair was removed from the dorsum of the mouse and cleaned. Five μg of S1 or S2 protein was mixed with 2 μg LPS immediately before subcutaneous deposition. Mice were anesthetized with isoflurane (Baxter, Deerfield, IL, USA) and preparations were injected subcutaneously either in the left or the right side of the dorsum. In one group of animals, to study the effect of polymyxin B on S2-induced NF-κB activation, 5 μg of S2 was mixed with 1 mg of polymyxin B (Sigma) immediately before subcutaneous deposition. Animals were then immediately transferred to individually ventilated cages. An In Vivo Imaging System (IVIS Spectrum, PerkinElmer Life Sciences) was used for the longitudinal imaging of NF-κB activation. Mice were imaged at 1, 3, and 6 h after the subcutaneous deposition. Fifteen minutes before the IVIS imaging, mice were intraperitoneally given 100 μl of D-luciferin (150 mg/kg body weight). Bioluminescence signals from the mice were detected and quantified using Living Image 4.0 Software (PerkinElmer Life Sciences).

## Data availability

The data supporting the findings of the study are available in the article and its Supporting Information or available upon request from the corresponding authors.

