## Supplementary Material for "SARS-CoV-2 spike protein as a bacterial lipopolysaccharide delivery system in an overzealous inflammatory cascade"

A

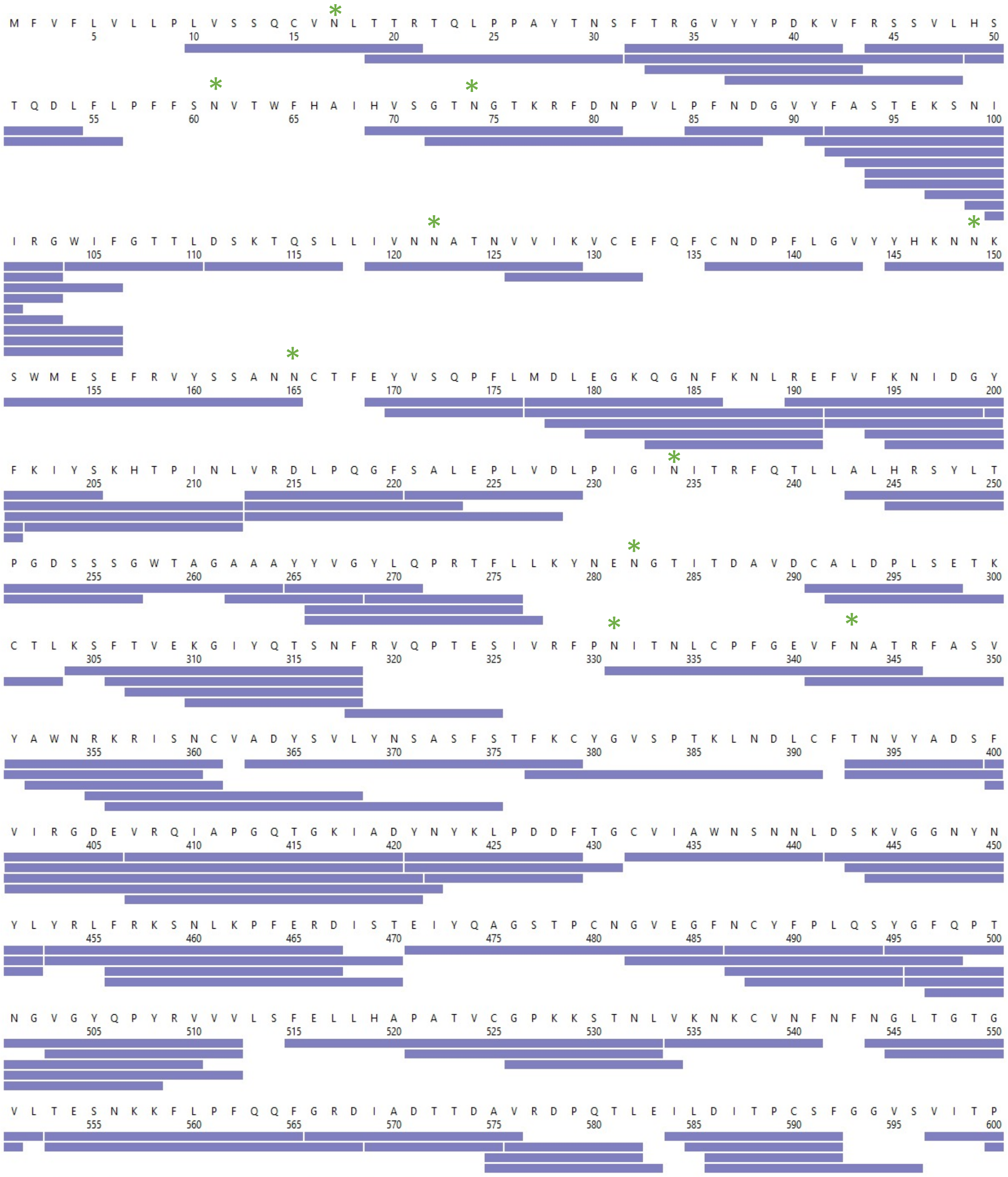

M F V F L V L L P L V S S Q C V N L T T R T Q L P P A Y T N S F T R G V Y Y P D K V F R S S V L H S
5 10 15 20 25 30 35 40 45 50
T Q D L F L P F F S N V T W F H A I H V S G T N G T K R F D N P V L P F N D G V Y F A S T E K S N I
55 60 65 70 75 80 85 90 95 100
I R G W I F G T T L D S K T Q S L L I V N N A T N V V I K V C E F Q F C N D P F L G V Y Y H K N N K
105 110 115 120 125 130 135 140 145 150
S W M E S E F R V Y S S A N N C T F E Y V S Q P F L M D L E G K Q G N F K N L R E F V F K N I D G Y
155 160 165 170 175 180 185 190 195 200
F K I Y S K H T P I N L V R D L P Q G F S A L E P L V D L P I G I N I T R F Q T L L A L H R S Y L T
205 210 215 220 225 230 235 240 245 250
P G D S S S G W T A G A A A Y Y V G Y L Q P R T F L L K Y N E N G T I T D A V D C A L D P L S E T K
255 260 265 270 275 280 285 290 295 300
C T L K S F T V E K G I Y Q T S N F R V Q P T E S I V R F P N I T N L C P F G E V F N A T R F A S V
305 310 315 320 325 330 335 340 345 350
Y A W N R K R I S N C V A D Y S V L Y N S A S F S T F K C Y G V S P T K L N D L C F T N V Y A D S F
355 360 365 370 375 380 385 390 395 400
V I R G D E V R Q I A P G Q T G K I A D Y N Y K L P D D F T G C V I A W N S N N L D S K V G G N Y N
405 410 415 420 425 430 435 440 445 450
Y L Y R L F R K S N L K P F E R D I S T E I Y Q A G S T P C N G V E G F N C Y F P L Q S Y G F Q P T
455 460 465 470 475 480 485 490 495 500
N G V G Y Q P Y R V V V L S F E L L H A P A T V C G P K K S T N L V K N K C V N F N F N G L T G T G
505 510 515 520 525 530 535 540 545 550
V L T E S N K K F L P F Q Q F G R D I A D T T D A V R D P Q T L E I L D I T P C S F G G V S V I T P
555 560 565 570 575 580 585 590 595 600

**B**

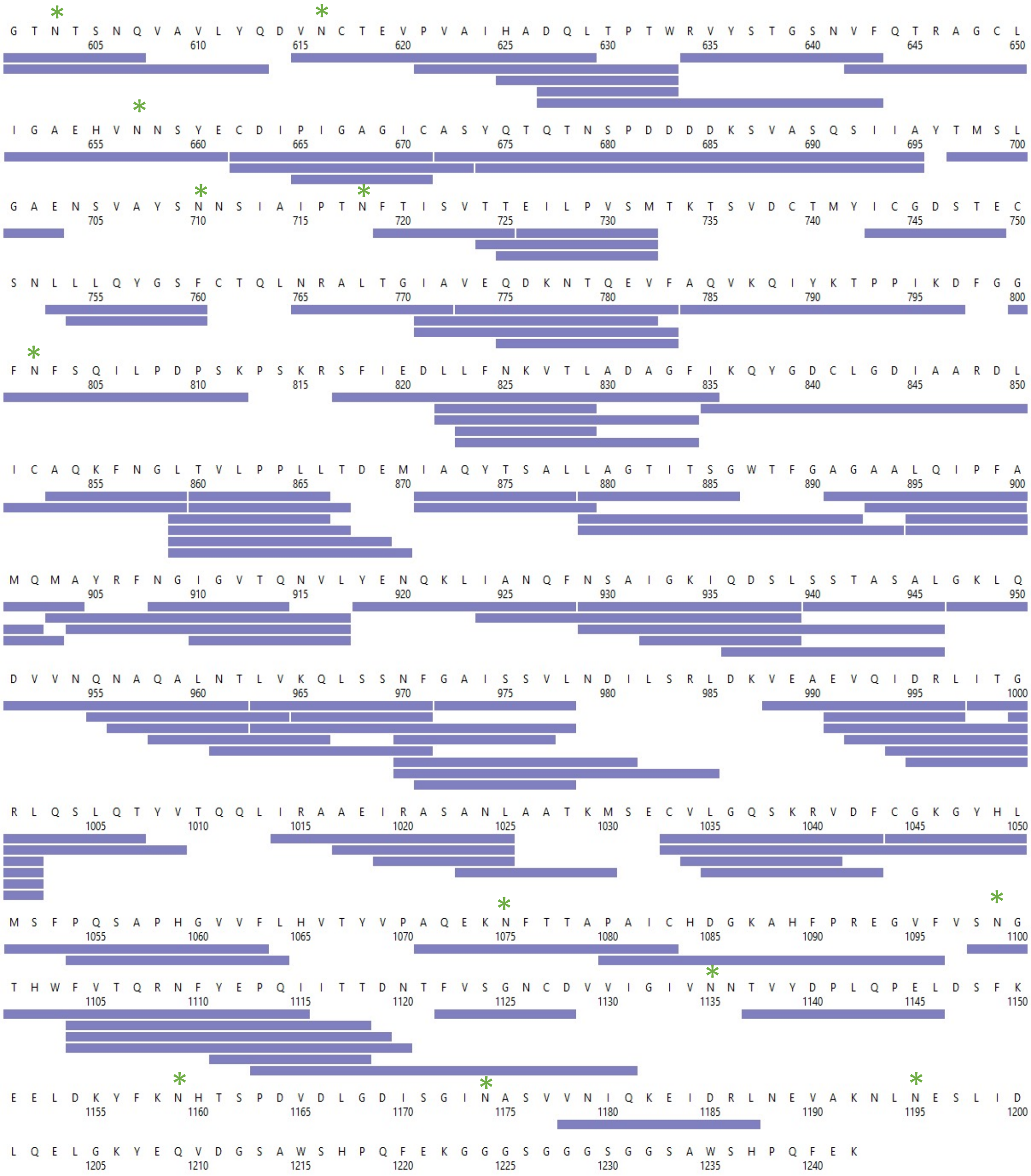
G T N T S N Q V A V L Y Q D V N C T E V P V A I H A D Q L T P T W R V Y S T G S N V F Q T R A G C L
605 610 615 620 625 630 635 640 645 650
I G A E H V N N S Y E C D I P I G A G I C A S Y Q T Q T N S P D D D D K S V A S Q S I I A Y T M S L
655 660 665 670 675 680 685 690 695 700
G A E N S V A Y S N N S I A I P T N F T I S V T T E I L P V S M T K T S V D C T M Y I C G D S T E C
705 710 715 720 725 730 735 740 745 750
S N L L L Q Y G S F C T Q L N R A L T G I A V E Q D K N T Q E V F A Q V K Q I Y K T P P I K D F G G
755 760 765 770 775 780 785 790 795 800
F N F S Q I L P D P S K P S K R S F I E D L L F N K V T L A D A G F I K Q Y G D C L G D I A A R D L
805 810 815 820 825 830 835 840 845 850
I C A Q K F N G L T V L P P L L T D E M I A Q Y T S A L L A G T I T S G W T F G A G A A L Q I P F A
855 860 865 870 875 880 885 890 895 900
M Q M A Y R F N G I G V T Q N V L Y E N Q K L I A N Q F N S A I G K I Q D S L S S T A S A L G K L Q
905 910 915 920 925 930 935 940 945 950
D V V N Q N A Q A L N T L V K Q L S S N F G A I S S V L N D I L S R L D K V E A E V Q I D R L I T G
955 960 965 970 975 980 985 990 995 1000
R L Q S L Q T Y V T Q Q L I R A A E I R A S A N L A A T K M S E C V L G Q S K R V D F C G K G Y H L
1005 1010 1015 1020 1025 1030 1035 1040 1045 1050
M S F P Q S A P H G V V F L H V T Y V P A Q E K N F T T A P A I C H D G K A H F P R E G V F V S N G
1055 1060 1065 1070 1075 1080 1085 1090 1095 1100
T H W F V T Q R N F Y E P Q I I T T D N T F V S G N C D V V I G I V N N T V Y D P L Q P E L D S F K
1105 1110 1115 1120 1125 1130 1135 1140 1145 1150
E E L D K Y F K N H T S P D V D L G D I S G I N A S V V N I Q K E I D R L N E V A K N L N E S L I D
1155 1160 1165 1170 1175 1180 1185 1190 1195 1200
L Q E L G K Y E Q V D G S A W S H P Q F E K G G G S G G G S G G S A W S H P Q F E K
1205 1210 1215 1220 1225 1230 1235 1240

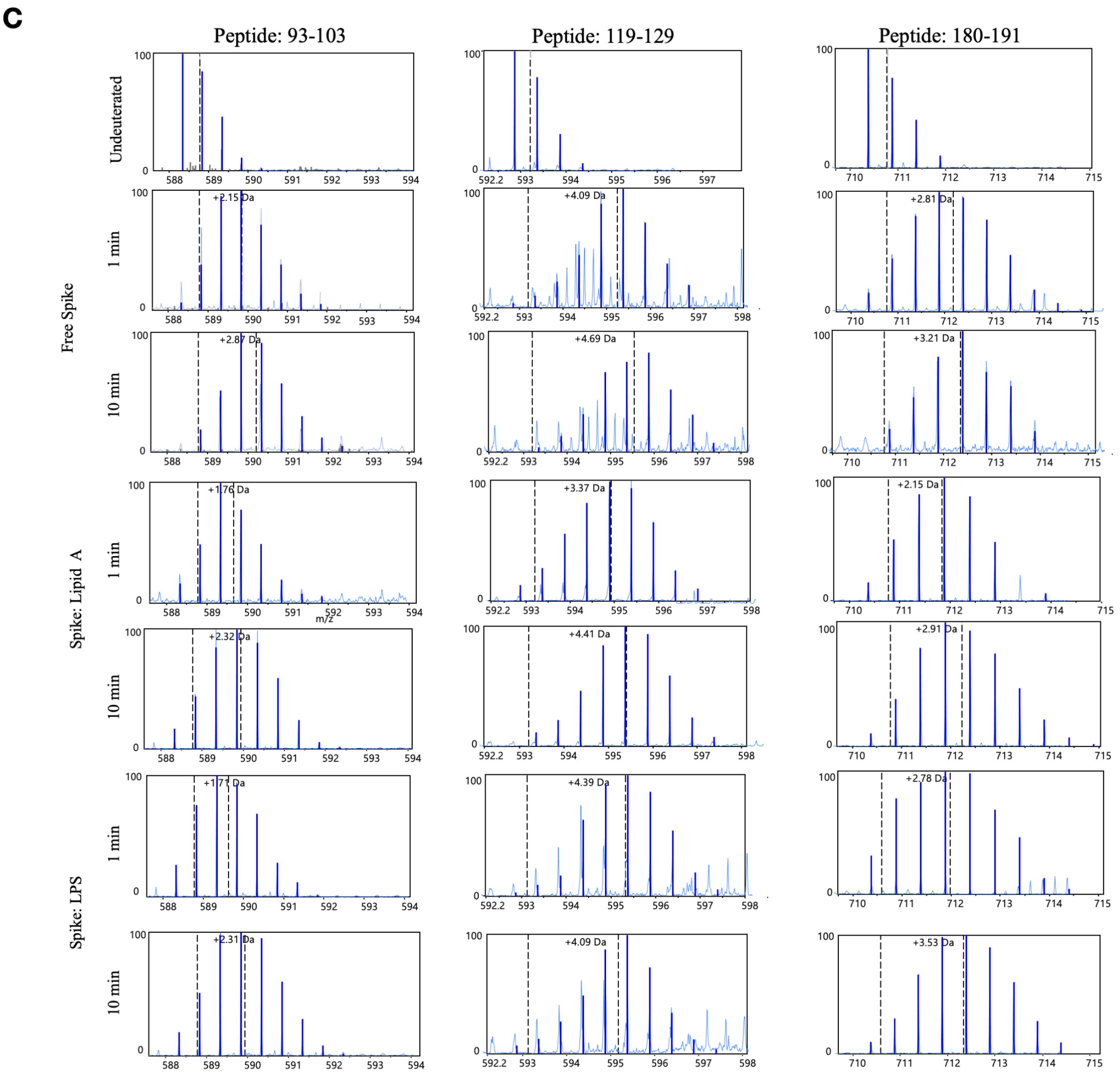

c
Peptide: 93-103
Peptide: 119-129
Peptide: 180-191
Undeuterated
Free Spike
Spike: Lipid A
Spike: LPS
1 min
10 min
+2.15 Da
+2.87 Da
+1.76 Da
+2.32 Da
+1.77 Da
+2.31 Da
+4.09 Da
+4.69 Da
+3.37 Da
+4.41 Da
+4.39 Da
+4.09 Da
+2.81 Da
+3.21 Da
+2.15 Da
+2.91 Da
+2.78 Da
+3.53 Da
m/z

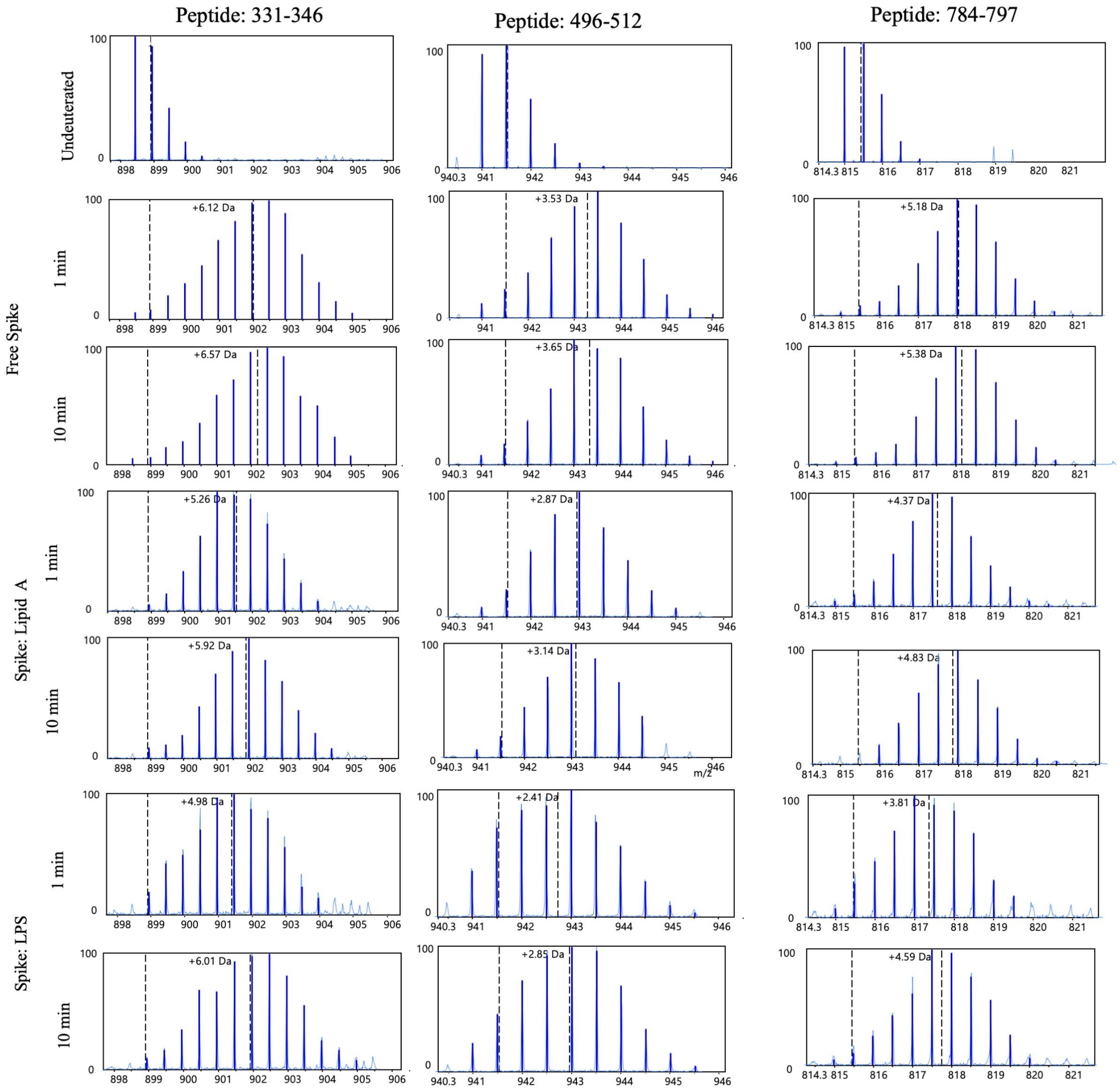

Peptide: 331-346
Peptide: 496-512
Peptide: 784-797
Undeuterated
Free Spike
1 min
10 min
Spike: Lipid A
1 min
10 min
Spike: LPS
1 min
10 min
+6.12 Da
+3.53 Da
+5.18 Da
+6.57 Da
+3.65 Da
+5.38 Da
+5.26 Da
+2.87 Da
+4.37 Da
+5.92 Da
+3.14 Da
+4.83 Da
+4.98 Da
+2.41 Da
+3.81 Da
+6.01 Da
+2.85 Da
+4.59 Da
m/z

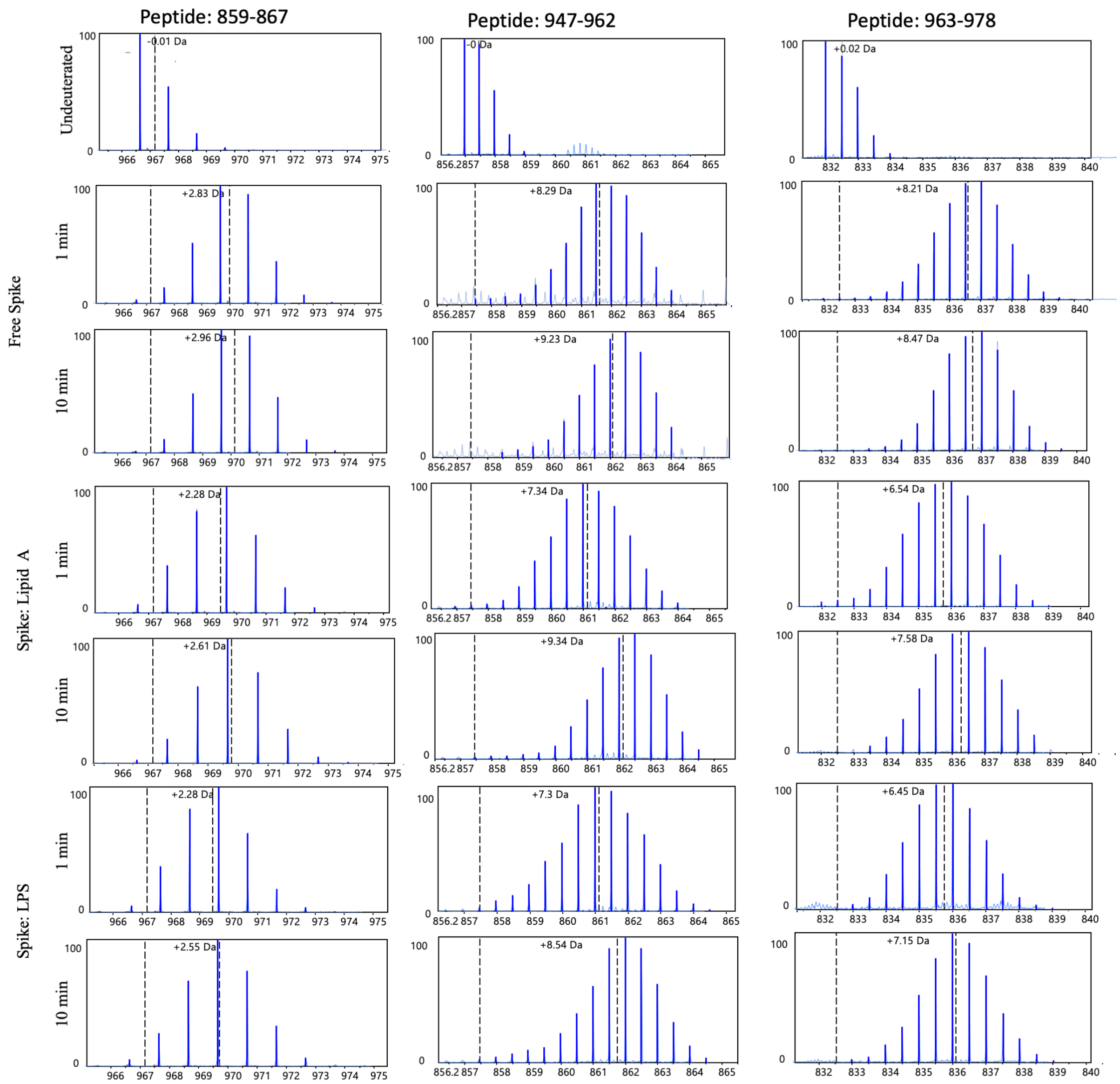


**Figure S1: Primary sequence coverage map of pepsin proteolyzed peptides of S protein.** Coverage map showing peptides spanning ~82% of the S protein: (A) 1 – 600 and (B) 601 – 1240 and glycosylation sites are indicated by asterisks (*) and peptide coverage for C-terminal twin strep-tag is not shown. (C) Mass spectra of representative peptides from NTD, RBD and S2 domain of Spike protein showing protection in the presence of lipid A and LPS at deuterium labelling times of 1 and 10 min. The dotted line highlights the centroids of undeuterated and deuterium labelled peptide mass spectra and difference between centroids shows the increases in mass after respective deuterium labelling times of each peptide.

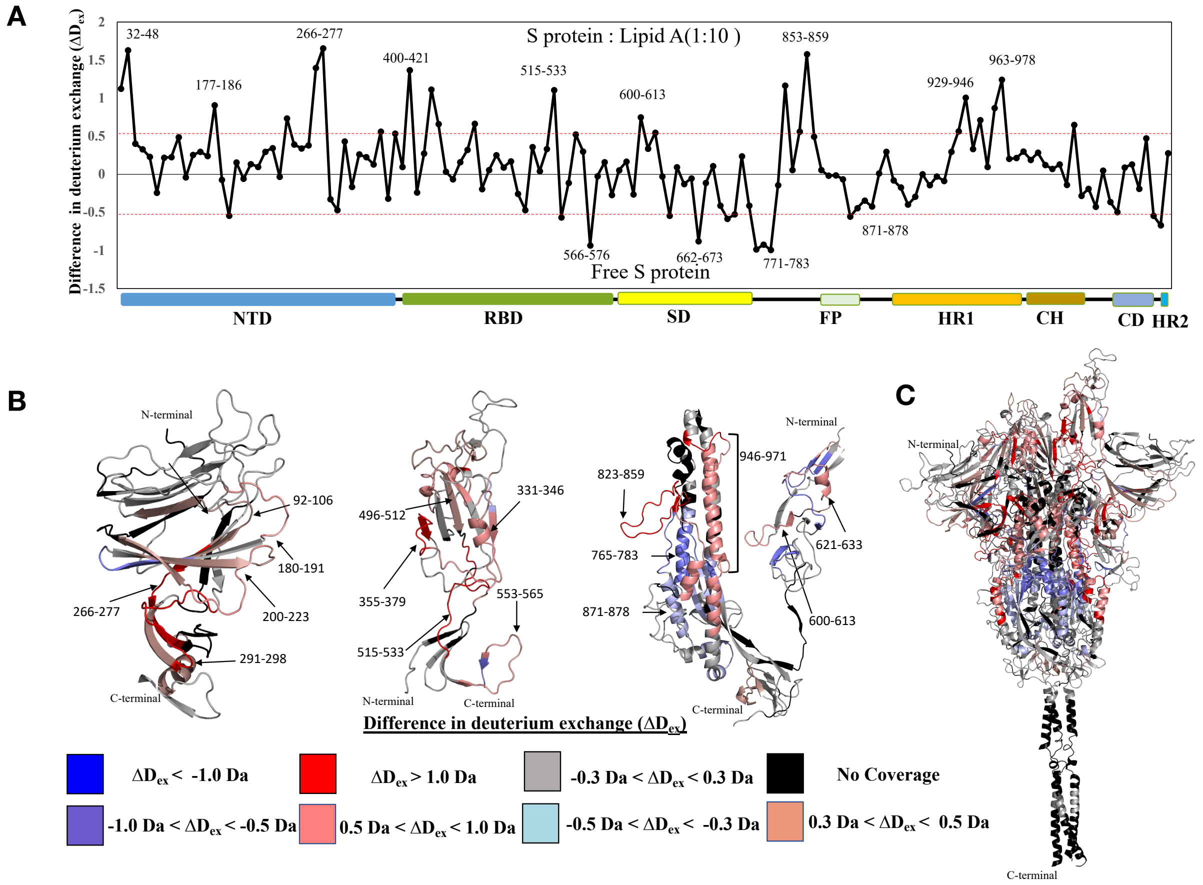


**Figure S2: Effect of Lipid A binding on S protein at 100 min deuterium labelling time.** (A) Plot showing differences in deuterium exchange ($\Delta D_{ex}$) between S protein: lipid A state and free S protein state at deuterium labelling times t = 100 min . Pepsin proteolyzed peptide fragments are grouped according to the S protein domain organisation from N to C-termini and plotted along X-axis and its corresponding difference in deuterium exchange values ($\Delta D_{ex}$) along Y-axis. A difference cut-off of ±0.5 Da is the significance threshold indicated with red dotted line. (B) Differences in deuterium exchange values at labelling times 1 and 10 min for S protein domains NTD (i), RBD (ii) and S2 domain (iii) are mapped on to the respective structures. (C) Differences in deuterium exchange values at labelling times t= 1 min and 10 min are mapped on to the full-length S protein (14-1208) structure. Deuterium exchange differences are colour coded as per key.

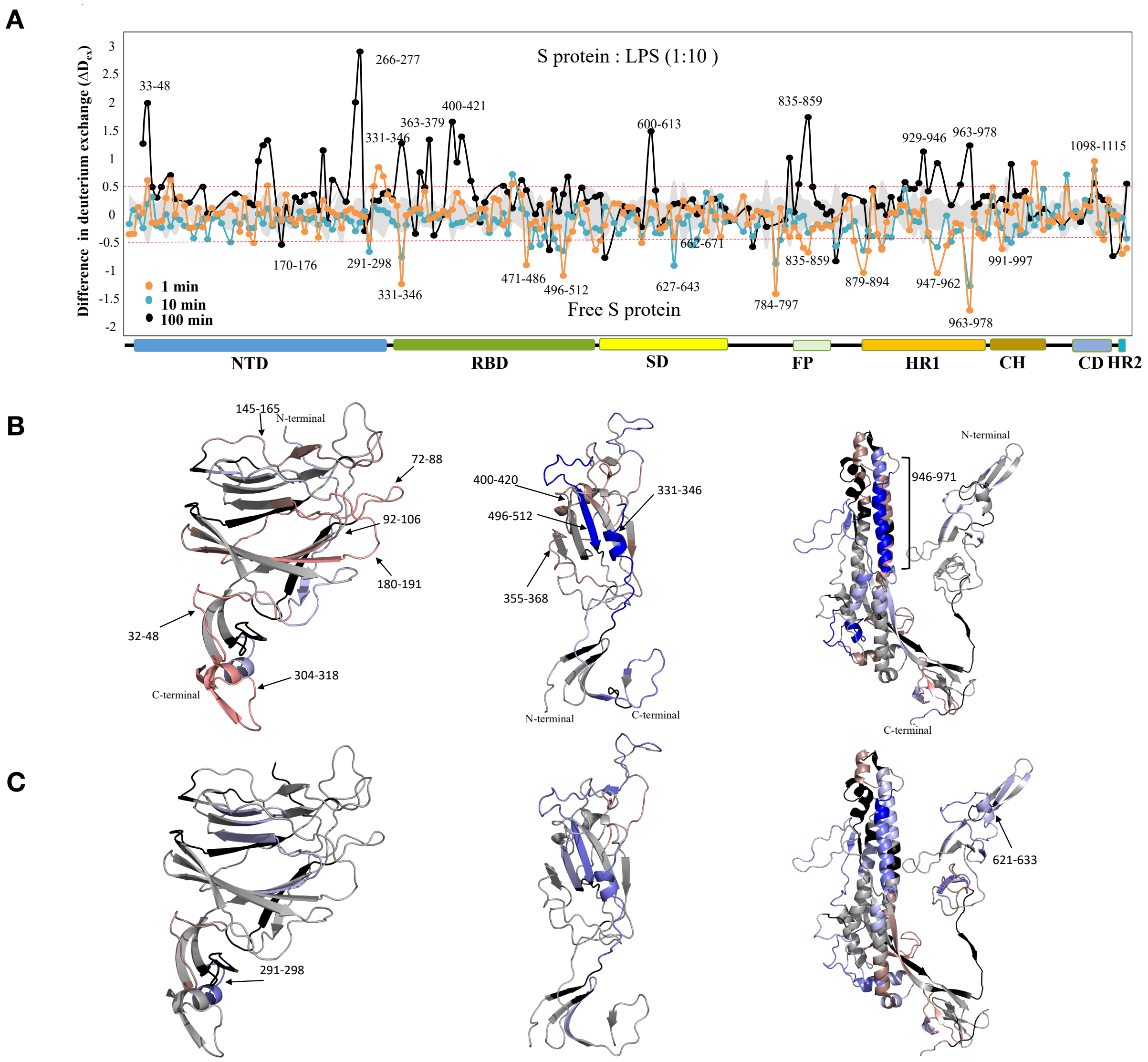

A
S protein : LPS (1:10 )
Difference in deuterium exchange (ΔDex)
33-48
266-277
400-421
363-379
331-346
600-613
835-859
929-946
963-978
1098-1115
170-176
291-298
331-346
471-486
496-512
627-643
662-671
784-797
835-859
879-894
947-962
963-978
991-997
1 min
10 min
100 min
Free S protein
NTD
RBD
SD
FP
HR1
CH
CD
HR2
B
145-165
N-terminal
72-88
92-106
180-191
32-48
304-318
C-terminal
400-420
496-512
331-346
355-368
N-terminal
C-terminal
946-971
N-terminal
C-terminal
C
291-298
621-633

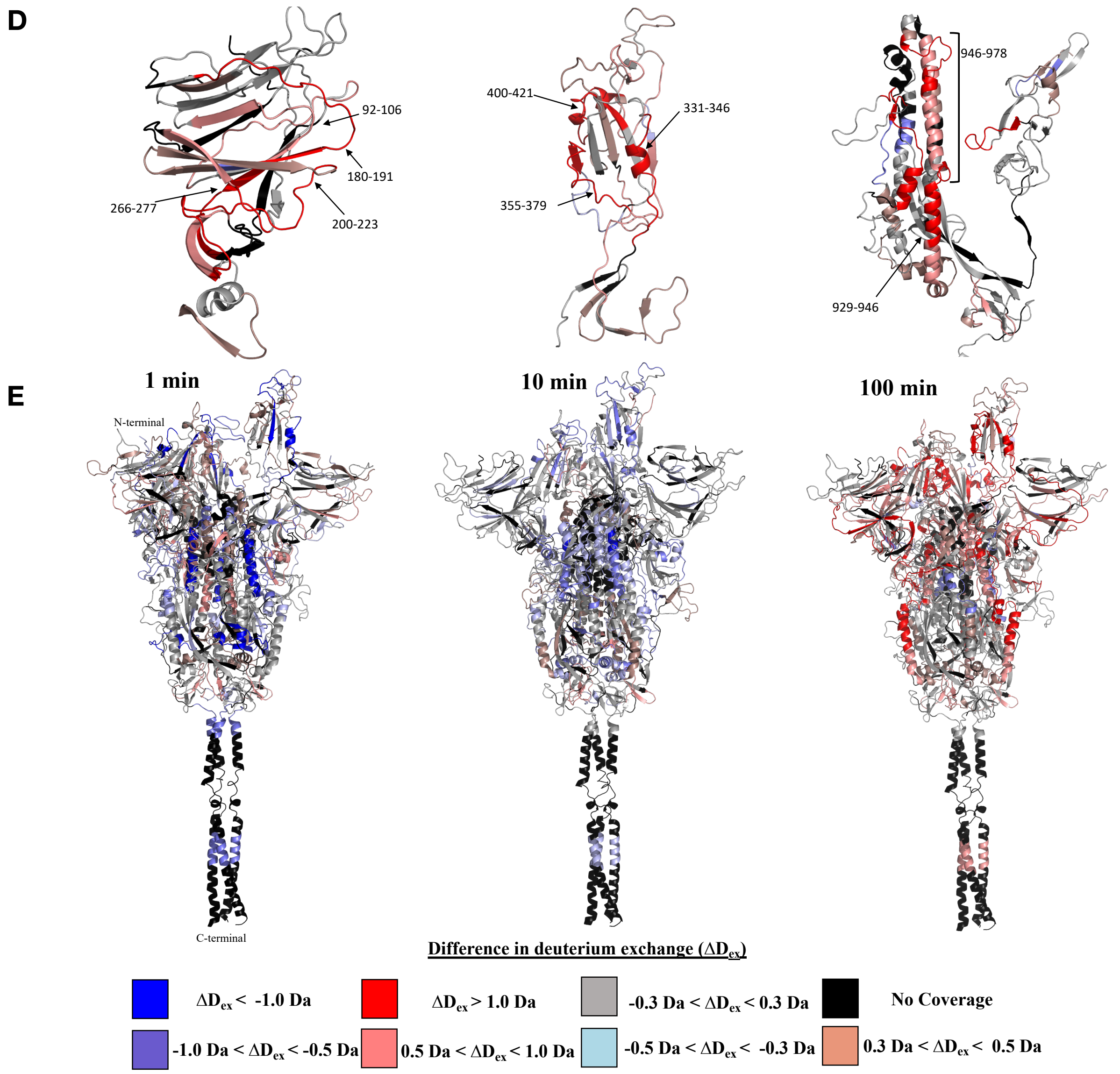


**Figure S3: Effect of LPS binding on Spike protein.** (A) Plot showing differences in deuterium exchange ($\Delta D_{ex}$) between S protein: lipid A state and free S protein state at deuterium labelling times t = 1 min, 10 min and 100 min . Pepsin proteolyzed peptide fragments are grouped according to the S protein domain organisation from N to C-termini and plotted along X-axis and its corresponding difference in deuterium exchange values ($\Delta D_{ex}$) along Y-axis. A difference cut-off of ±0.5 Da is the significance threshold indicated with red dotted line. (B-D) Differences in deuterium exchange ($\Delta D_{ex}$) values at labelling times 1 min, 10 min and 100 min for S protein domains NTD (i), RBD (ii) and S2 domain (iii) are mapped on to the respective structures. (C) Differences in deuterium exchange values at labelling times t= 1 min, 10 min and 100 min are mapped on to the full-length S protein (14-1208) structure. Deuterium exchange differences are colour coded as per key.

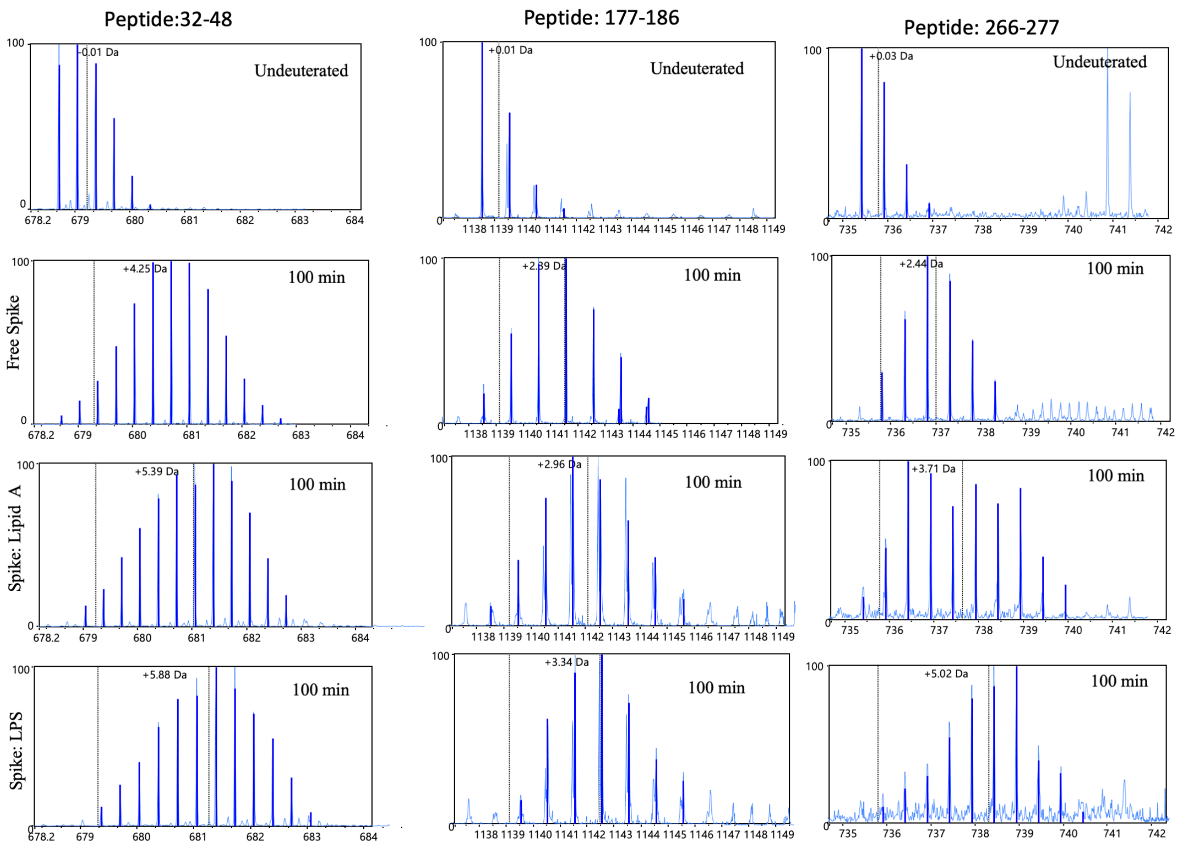

Peptide:32-48
Peptide: 177-186
Peptide: 266-277
Undeuterated
100 min
Free Spike
Spike: Lipid A
Spike: LPS
+0.01 Da
+0.03 Da
+4.25 Da
+2.89 Da
+2.44 Da
+5.39 Da
+2.96 Da
+3.71 Da
+5.88 Da
+3.34 Da
+5.02 Da

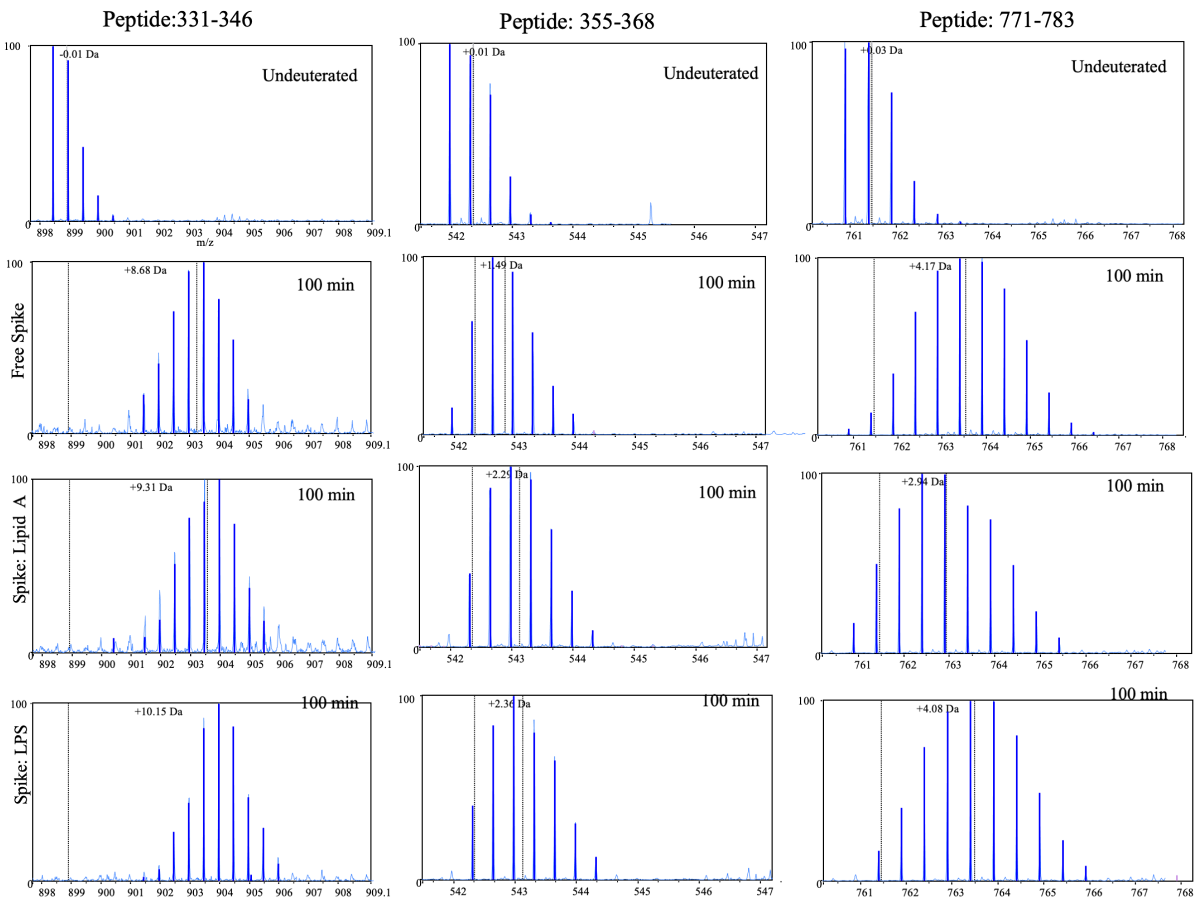

Peptide:331-346
Peptide: 355-368
Peptide: 771-783
Undeuterated
100 min
Free Spike
Spike: Lipid A
Spike: LPS
-0.01 Da
+0.01 Da
+0.03 Da
+8.68 Da
+1.49 Da
+4.17 Da
+9.31 Da
+2.29 Da
+2.94 Da
+10.15 Da
+2.36 Da
+4.08 Da
m/z

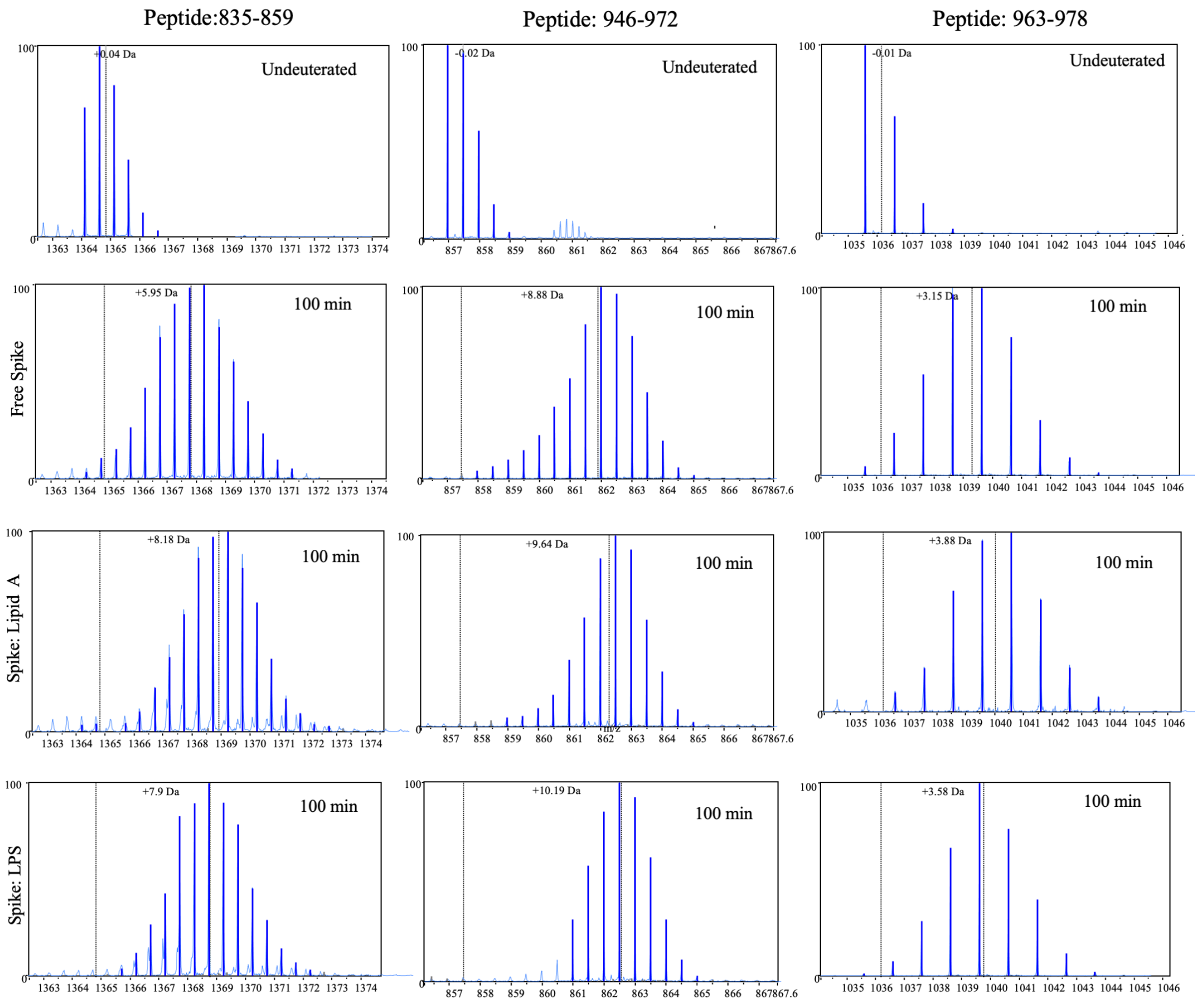


**Figure S4: Comparing mass spectra of representative peptides showing increases in deuterium exchange at 100 min labelling from NTD, RBD and S2 domains.** Increases in deuterium exchange at longer time points indicate destabilisation of the Spike trimer at the inter-protomer interface due to insertion of lipid tails. Also, flexible lipid tails binding at the cavities at the NTD and RBD pockets may induce destabilisation of local secondary structure. Peptide 266-277 spans across NTD (266-271) and S2 (272-277) binding pockets, and shows bimodal spectra. This may result from a cumulative effect of lipid A binding, wherein the low and high exchanging populations represent protection from deuterium exchange due to lipid A binding at NTD and destabilization caused by multiple lipid A molecules binding at the S2 pocket. Interestingly, bimodal spectra were not observed in the case of the LPS bound state indicating a uniform effect, i.e., destabilization by bulky LPS binding at both NTD and S2 pockets.

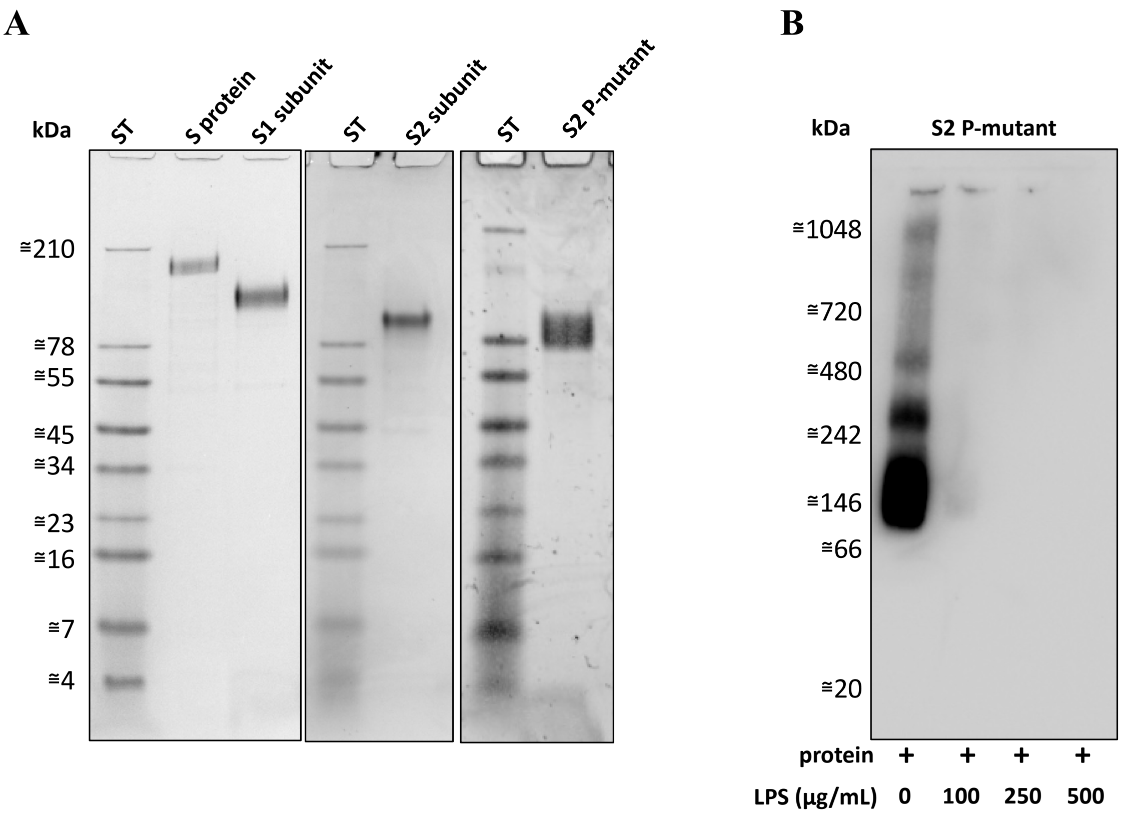


**Figure S5: Evaluation of SARS-CoV-2 S protein and its subunits on SDS- and BN-PAGE**. (A) 2 µg of S protein, S1, S2 or S2 P-mutant (S2 subunits with proline substitutions (F817P, A892P, A899P, A942P, K986P, V987P)) subunits, were separated by SDS-PAGE under reducing conditions, and then stained with Coomassie. One representative image of two independent experiments is shown (n=2). (B) S2 P-mutant was incubated with (0–500 µg/ml) LPS, separated by BN-PAGE and detected by western blotting. One representative image of three independent experiments is shown (n=3).

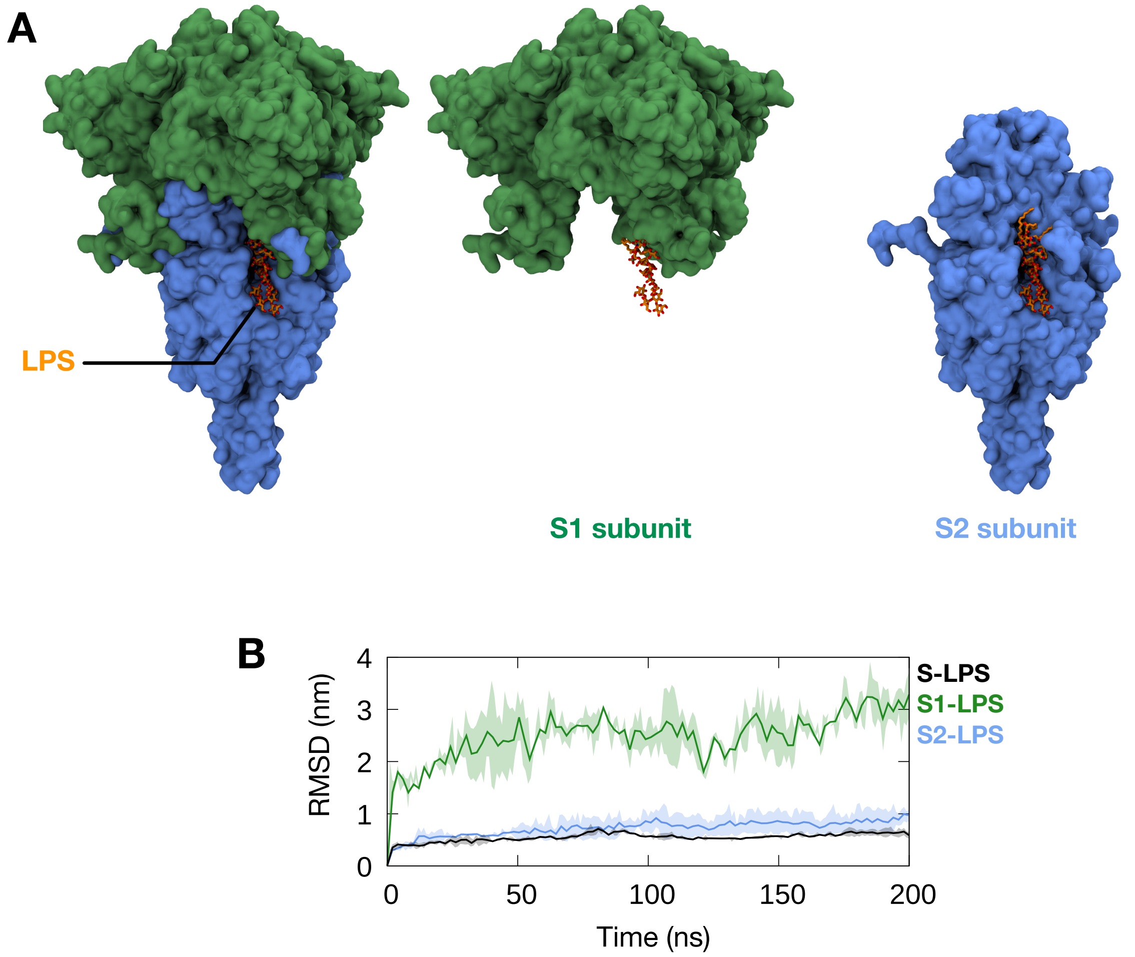


**Figure S6: Simulations of LPS bound to S2 binding pocket in full-length spike or truncated S1 or S2 systems.** (A) Initial snapshots of simulations. LPS is shown in orange and stick representation bound to the S2 binding site, with the S1 subunit in green and S2 subunit in cyan, both depicted in surface representation. (B) The root mean square deviation (RMSD) of LPS throughout the simulations, after performing a least-squares fit to the protein backbone, compared to simulations of LPS bound to the whole S ECD (black). Thick line shows average values for three independent simulations and the shaded areas indicate standard deviations.

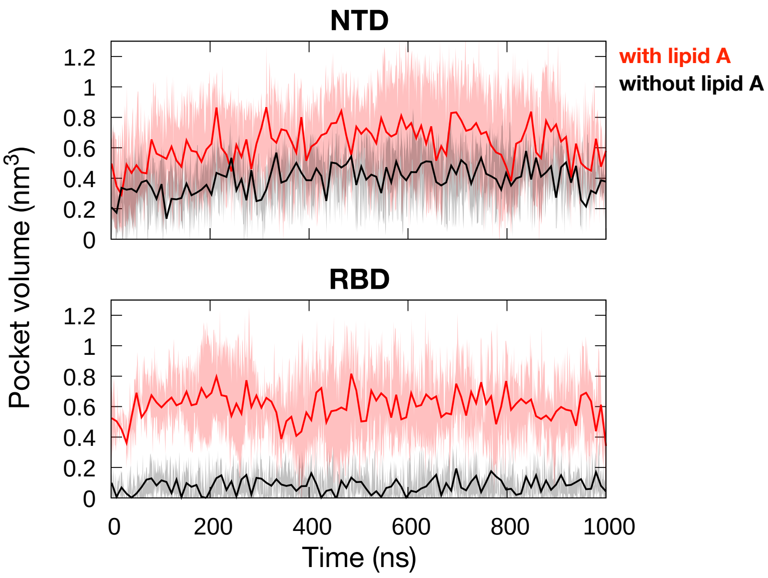


**Figure S7: The volume of NTD and RBD pockets when bound to lipid A.** The volume of hydrophobic pockets throughout three independent simulations with and without lipid A. Thick lines show average values, and the shaded areas indicate standard deviations.

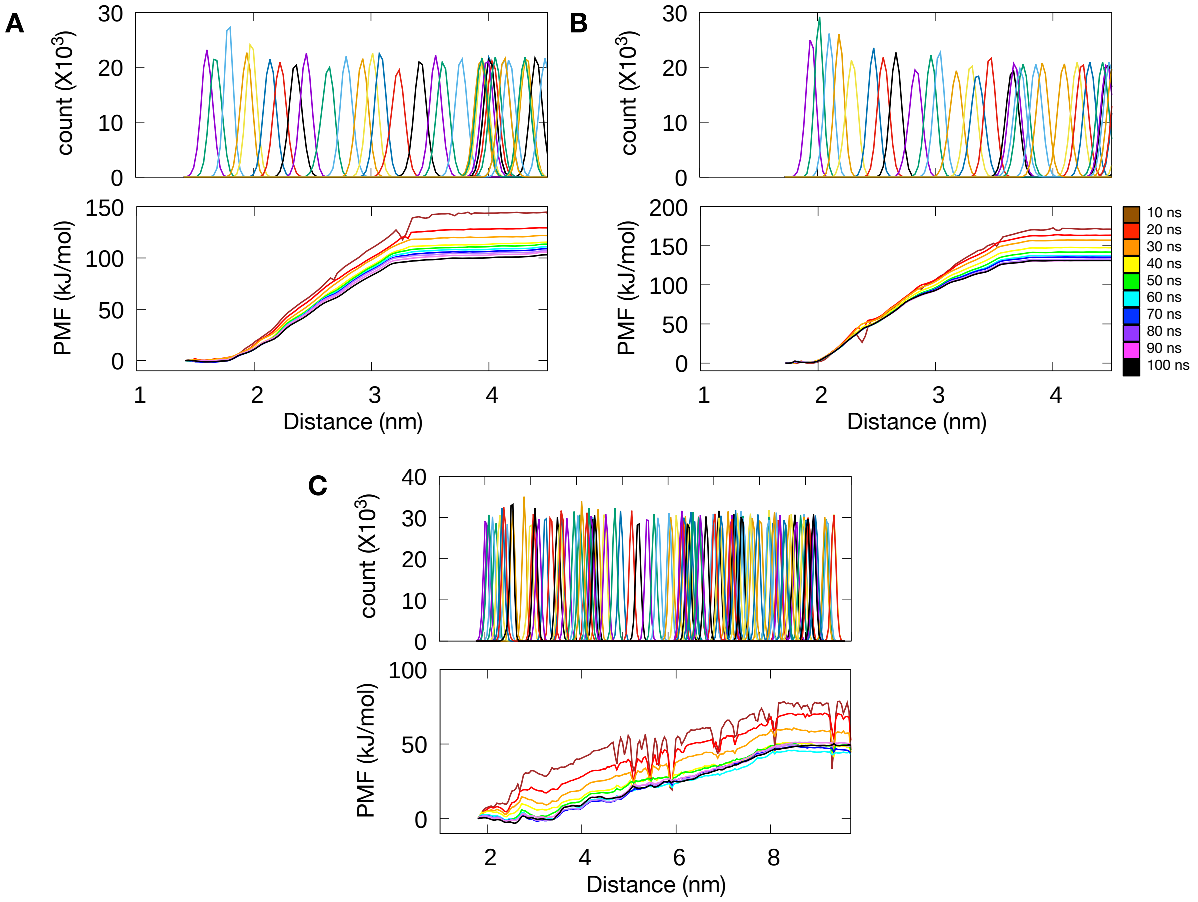


**Figure S8: Sampling and convergence analyses of PMF calculations.** (A) (Top) Histogram overlap from all umbrella sampling windows of lipid A dissociation from the NTD pocket. (Bottom) PMF profiles generated with increasing lengths of simulation sampling. (B and C) Similar analyses performed on PMF calculations for lipid A dissociation from the RBD pocket and the S2 pocket, respectively. For the NTD pocket, the PMF profiles converged after 40 ns, while for the RBD and S2 pockets, the PMF profiles converged after 60 ns.

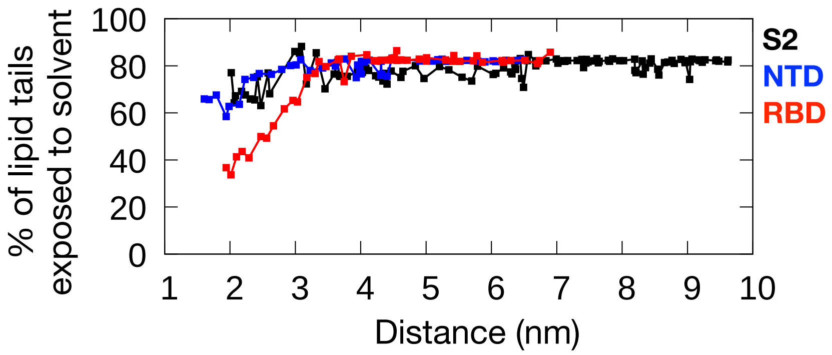


**Figure S9: Percentage of lipid tails atoms exposed to water during umbrella sampling simulations.** Solvent exposed atoms are defined as those found within 0.4 nm of any water molecule. The values are averaged over the converged portion of each umbrella sampling window.

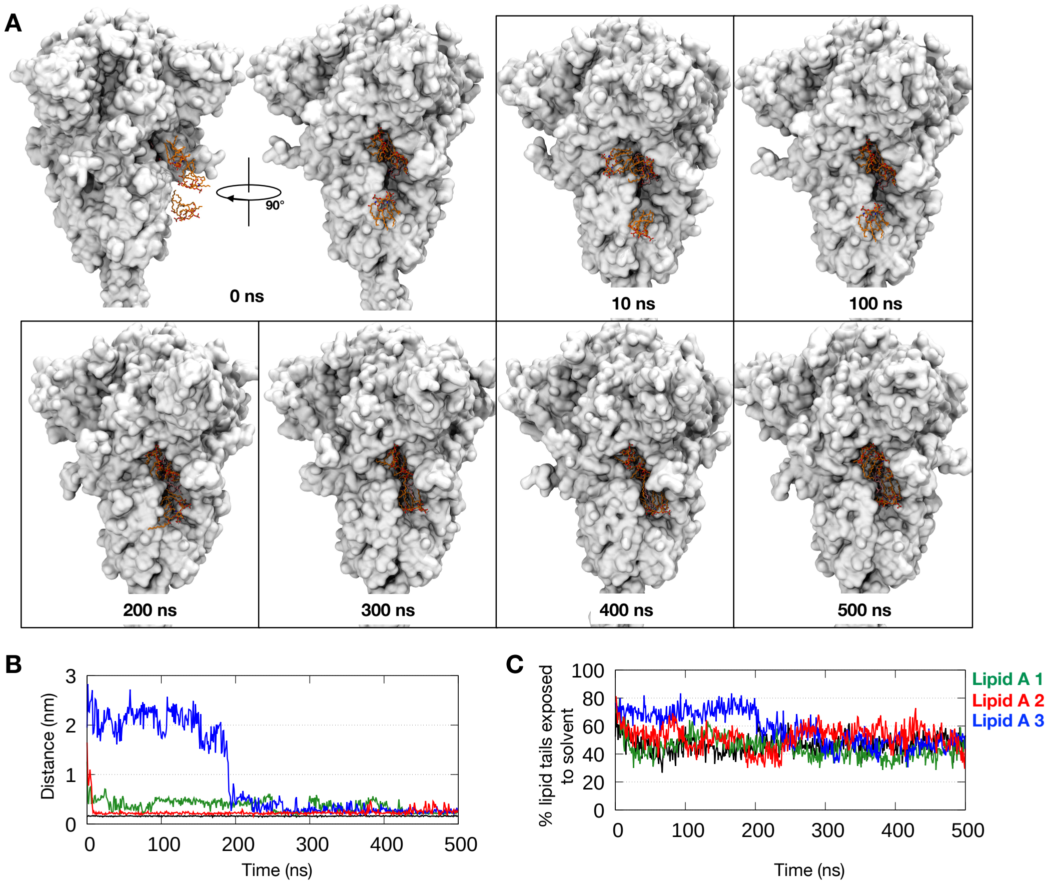


**Figure S10: Unbiased simulation of multiple lipid A molecules binding to S2 binding site.** (A) Three lipid A molecules were placed outside of the S2 pocket with their centers of mass located at least 2 nm from an already bound lipid A molecule. Snapshots were taken at different time points during the simulation, with lipid A shown in orange and stick representation and protein shown in white surface representation. (B) Minimum distance between three unbound lipid A molecules (green, red and blue) from protein residues forming the binding site as described in our previous study.[1] The distance from an already bound lipid A is shown in black as reference. (C) The percentage of lipid tail atoms from each lipid A molecule that were exposed to solvent during the simulation. Solvent exposed atoms are defined as those found within 0.4 nm of any water molecule. Black line indicates data for the lipid A molecule that was already bound at the beginning of the simulation.

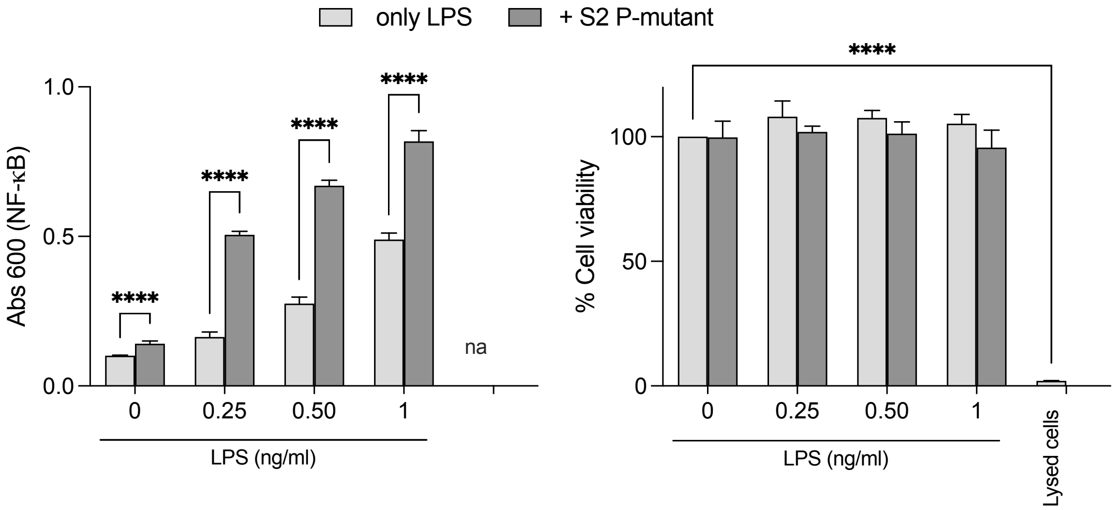


**Figure S11: S2 P-mutant boosts the pro-inflammatory response to LPS in human monocytes.** NF-κB activation (left panel) and cell viability (right panel) were measured in THP-1-XBlue-CD14 cells stimulated with increasing doses of LPS (0 to 0.25 ng/ml), with or w/o 5 nM of S2 P-mutant. Lysed cells were used as negative control for cell viability. Data are presented as mean ± SD of three independent experiments performed in triplicate (n=3). ****$P \leq 0.0001$, determined using two-way ANOVA with Tukey's multiple comparisons test.

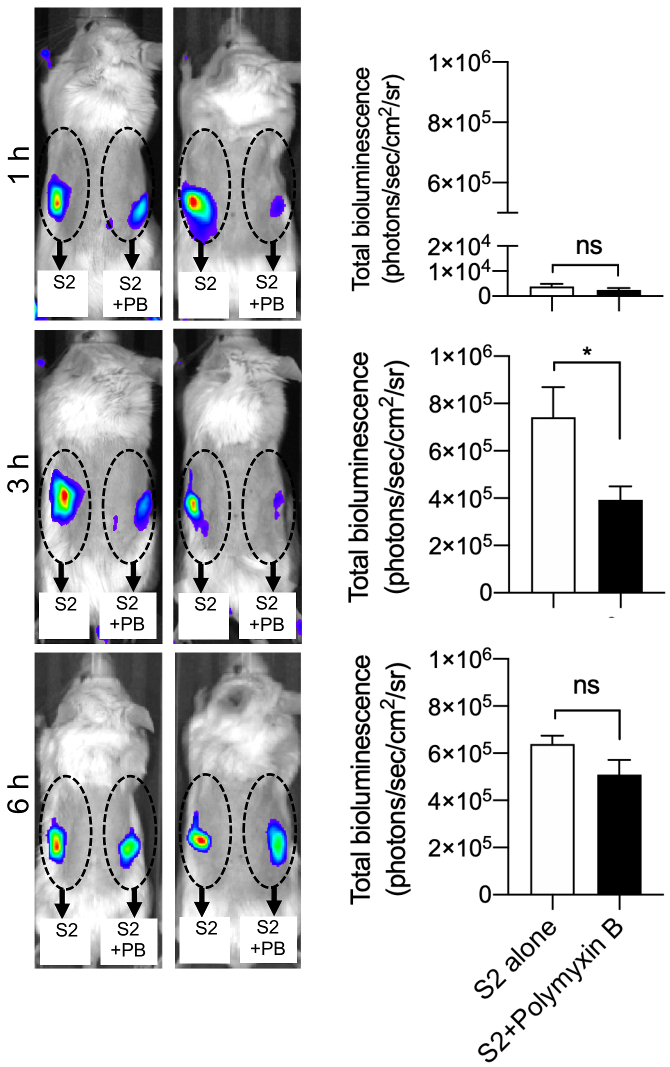


**Figure S12: Effect of polymyxin B on S2-induced NF-κB activation**. Bioimaging of inflammation in NF-κB reporter mice. S2 alone or in combination with polymyxin B (PB) was subcutaneously deposited on the left and right side on the back of transgenic BALB/c Tg(NF-κB-RE-luc)-Xen reporter mice. *In vivo* imaging of NF-κB reporter gene expression was performed using the IVIS Spectrum system. Representative images show bioluminescence at 1, 3 and 6 h after subcutaneous deposition. Bar charts show bioluminescence emitted from these reporter mice. Areas of subcutaneous deposition and region of interest for bioluminescence analysis are depicted as dotted circles. Data are presented as the mean ± SEM ($n$ = 4). $P$ values were determined using a one-way ANOVA with Dunnett posttest. *$P \leq 0.05$; ns, not significant.

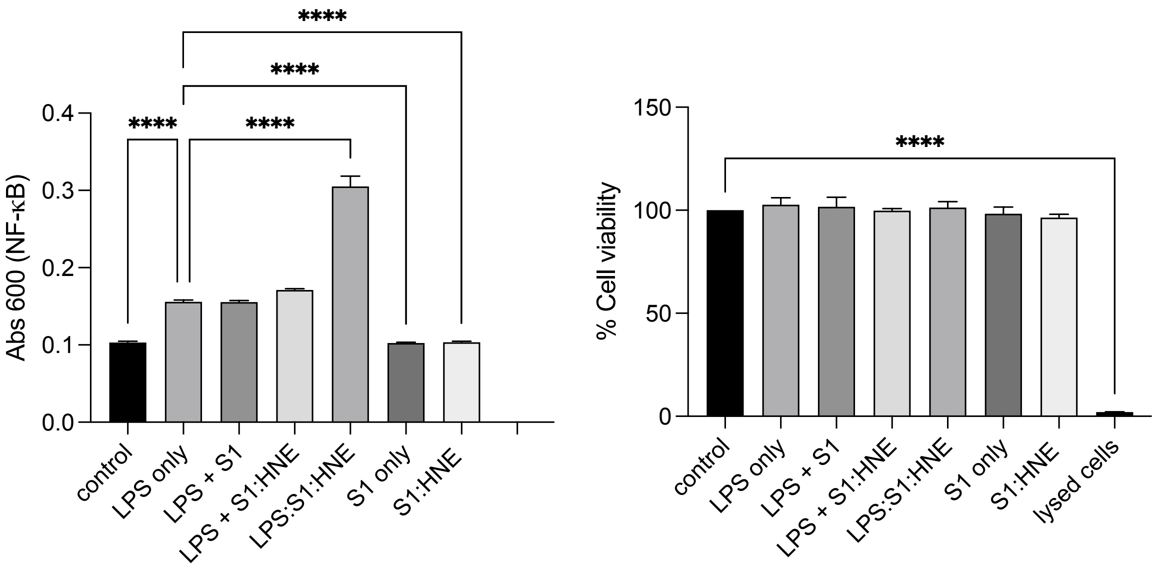


**Figure S13: S1 subunit boosts the pro-inflammatory response to LPS after digestion with human neutrophil elastase (HNE).** NF-κB activation (left panel) and cell viability (right panel) were measured in THP-1-XBlue-CD14 cells stimulated with 0.25 ng/ml of LPS alone, LPS and 5 nM of S1 (LPS +S1), LPS with digested S1 subunit with HNE (LPS + S1:HNE), S1 subunit digested with HNE in the presence of LPS (LPS:S1:HNE), intact or digested with HNE (S1:HNE) S1 subunit. Lysed cells were used as negative control for cell viability. Data are presented as mean ± SD of three independent experiments performed in triplicate (n=3). ****$P \leq 0.0001$, determined using two-way ANOVA with Tukey's multiple comparisons test.
